# Highly specific σ_2_R/TMEM97 ligand alleviates neuropathic pain and inhibits the integrated stress response

**DOI:** 10.1101/2023.04.11.536439

**Authors:** Muhammad Saad Yousuf, James J. Sahn, Hongfen Yang, Eric T. David, Stephanie Shiers, Marisol Mancilla Moreno, Jonathan Iketem, Danielle M. Royer, Chelsea D. Garcia, Jennifer Zhang, Veronica M. Hong, Subhaan M. Mian, Ayesha Ahmad, Benedict J. Kolber, Daniel J Liebl, Stephen F. Martin, Theodore J. Price

## Abstract

The Sigma 2 receptor (σ_2_R) was described pharmacologically more than three decades ago, but its molecular identity remained obscure until recently when it was identified as transmembrane protein 97 (TMEM97). We and others have shown that σ_2_R/TMEM97 ligands alleviate mechanical hypersensitivity in mouse neuropathic pain models with a time course wherein maximal anti-nociceptive effect is approximately 24 hours following dosing. We sought to understand this unique anti-neuropathic pain effect by addressing two key questions: do these σ_2_R/TMEM97 compounds act selectively via the receptor, and what is their downstream mechanism on nociceptive neurons? Using male and female conventional knockout (KO) mice for *Tmem97,* we find that a new σ_2_R/TMEM97 binding compound, FEM-1689, requires the presence of the gene to produce anti-nociception in the spared nerve injury model in mice. Using primary mouse dorsal root ganglion (DRG) neurons, we demonstrate that FEM-1689 inhibits the integrated stress response (ISR) and promotes neurite outgrowth via a σ_2_R/TMEM97-specific action. We extend the clinical translational value of these findings by showing that FEM-1689 reduces ISR and p-eIF2α levels in human sensory neurons and that it alleviates the pathogenic engagement of ISR by methylglyoxal. We also demonstrate that σ_2_R/TMEM97 is expressed in human nociceptors and satellite glial cells. These results validate σ_2_R/TMEM97 as a promising target for further development for the treatment of neuropathic pain.

**Significance Statement:** Neuropathic pain is a major medical problem that is poorly treated with existing therapeutics. Our findings demonstrate that targeting σ_2_R/TMEM97 with a newly described modulator reduces pain hypersensitivity in a mouse model with exquisite selectivity. We also identify integrated stress response (ISR) inhibition as a potential mechanism of action that links the receptor to cellular signaling events that have preclinical and clinical validation for pain relief. Our work suggests that σ_2_R/TMEM97 can be selectively engaged by specific small molecules to produce ISR inhibition in a subset of cells that are critical for neuropathic pain. σ_2_R/TMEM97-targeted therapeutics thus have the potential to offer effective pain relief without engagement of opioid receptors.

## Introduction

Neuropathic pain, which is caused by an injury or disease of the somatosensory nervous system, affects approximately 10% of the population and is the leading cause of high-impact chronic pain (1). Management of neuropathic pain is a major clinical challenge because available drugs not only have limited efficacy, but they also elicit serious side effects. There is a significant need for novel drugs that alleviate neuropathic pain through non-opioid and non-addicting mechanisms and have improved side effect profiles.

The sigma 2 receptor (σ_2_R) was identified in 2017 as transmembrane protein 97 (TMEM97) (2). We discovered that several small molecules that bind selectively to σ_2_R/TMEM97 produce strong and long-lasting anti-neuropathic pain effects from spared nerve injury (SNI) in mice (3), a finding that was independently replicated with structurally distinct molecules (4). Although the biological function of σ_2_R/TMEM97 is not well understood, it is a transmembrane protein that is associated with the endoplasmic reticulum (ER) and plays a role in calcium signaling (5, 6) and cholesterol trafficking and homeostasis (7–11). 20(*S*)-Hydroxycholesterol was recently identified as an endogenous ligand for σ_2_R/TMEM97 (12). The role of σ_2_R/TMEM97 in disease pathology has historically been focused on cancer (13), but it is also implicated in neurodegenerative diseases including Alzheimer’s disease (14–17) and Parkinson’s disease (18). Pharmacological targeting of σ_2_R/TMEM97 has neuroprotective effects in a number of models of neurodegenerative conditions, including traumatic brain injury (19), Huntington’s disease (20), and retinal ganglion cell degeneration (21).

The mechanism by which modulation of σ_2_R/TMEM97 alleviates neuropathic pain is not known. Given the localization of σ_2_R/TMEM97 at the ER, a primary hypothesis tested in our work is whether σ_2_R/TMEM97 targeting may reduce pain via interference with the integrated stress response (ISR), which includes ER stress. The ISR is an adaptive response to cellular stressors such as accumulation of misfolded proteins, lipid and oxidative stress, amino acid and heme deprivation, and viral infection (22, 23). A canonical signaling event associated with the ISR is phosphorylation of eukaryotic initiation factor 2α (eIF2α) in response to cellular stress conveyed by four kinases: protein kinase R (PKR), PKR-like ER kinase (10), heme-regulated inhibitor (HRI), and general control nonderepressible 2 (GCN2). Phosphorylation of eIF2α inhibits global protein synthesis and promotes the translation of mRNAs such as activated transcription factor 4 (ATF4). We and others have demonstrated that the induction of the ISR is associated with neuropathic pain caused by traumatic nerve injury (24, 25), metabolic disorders (26–28), and autoimmune disorders (29–31).

Another key question is whether the anti-neuropathic pain effects of σ_2_R/TMEM97 ligands are specifically due to their binding to σ_2_R/TMEM97 because such compounds can also have substantial activity at the sigma 1 receptor (σ_1_R), a receptor that also promotes anti-nociception in animal models (32–35). We used a knockout mouse of the *Tmem97* gene, and a new small molecule, FEM-1689, that has improved selectivity for σ_2_R/TMEM97 to test the hypothesis that σ_2_R/TMEM97 is causatively linked to anti-nociception in mouse neuropathic pain models. Indeed, the anti-nociceptive effect of FEM-1689 in the SNI model is completely absent in TMEM97KO mice. Our findings also show that FEM-1689 inhibits the ISR in a σ_2_R/TMEM97-dependent fashion in mouse and human DRG neurons. This work provides a strong mechanistic case for targeting of σ_2_R/TMEM97 as the basis of a new approach to treat neuropathic pain.

## Results

### TMEM97 mRNA is expressed in the human and mouse dorsal root ganglia

To assess whether σ_2_R/TMEM97 is expressed in human nociceptors, we performed RNAscope *in situ* hybridization using human dorsal root ganglia (DRG) obtained from organ donors. We found that *TMEM97* is expressed in all classes of human sensory neurons including *SCN10A* (Nav1.8)-positive, putative nociceptors (**Fig 1A-C**). Approximately 70% of all human DRG neurons expressed *SCN10A* transcripts, consistent with our previous observations (36), and all neurons evaluated expressed *TMEM97* mRNA (**Fig 1C**). Cell size distribution matrix of *TMEM97* expressing neurons (average = 74 µm) showed that *TMEM97* mRNA was not restricted to a subpopulation of neurons (**Fig 1D**). Upon further investigation, we found that *FABP7*-positive satellite glial cells also express *TMEM97* (**Fig 1E**). We identified and quantified *TMEM97*-expressing neuronal subpopulations in the human DRG using our previously published human DRG spatial RNA sequencing dataset (**Fig 1F**) (36). We validated our findings that *TMEM97* is expressed across all neuronal subtypes in the human DRG with particularly high expression in proenkephalin (PENK)-positive nociceptors and Aδ low-threshold mechanoreceptors (**Fig 1F**). Mouse DRG neurons and satellite glial cells express *Tmem97* mRNA (**Supp Fig 1A**) with a nearly identical pattern to what is seen in the human DRG. Moreover, previously published single-cell RNA sequencing data from mice shows that *Tmem97* is expressed in all DRG neuronal subtypes with particularly high expression in non-peptidergic neurons, Schwann cells, and satellite glial cells (37). The recently developed harmonized dataset of rodent, non-human primate, and human DRGs show that *TMEM97* is expressed in all sensory neuron types (**Supp Fig 1B**) (38).

**Figure 1.**
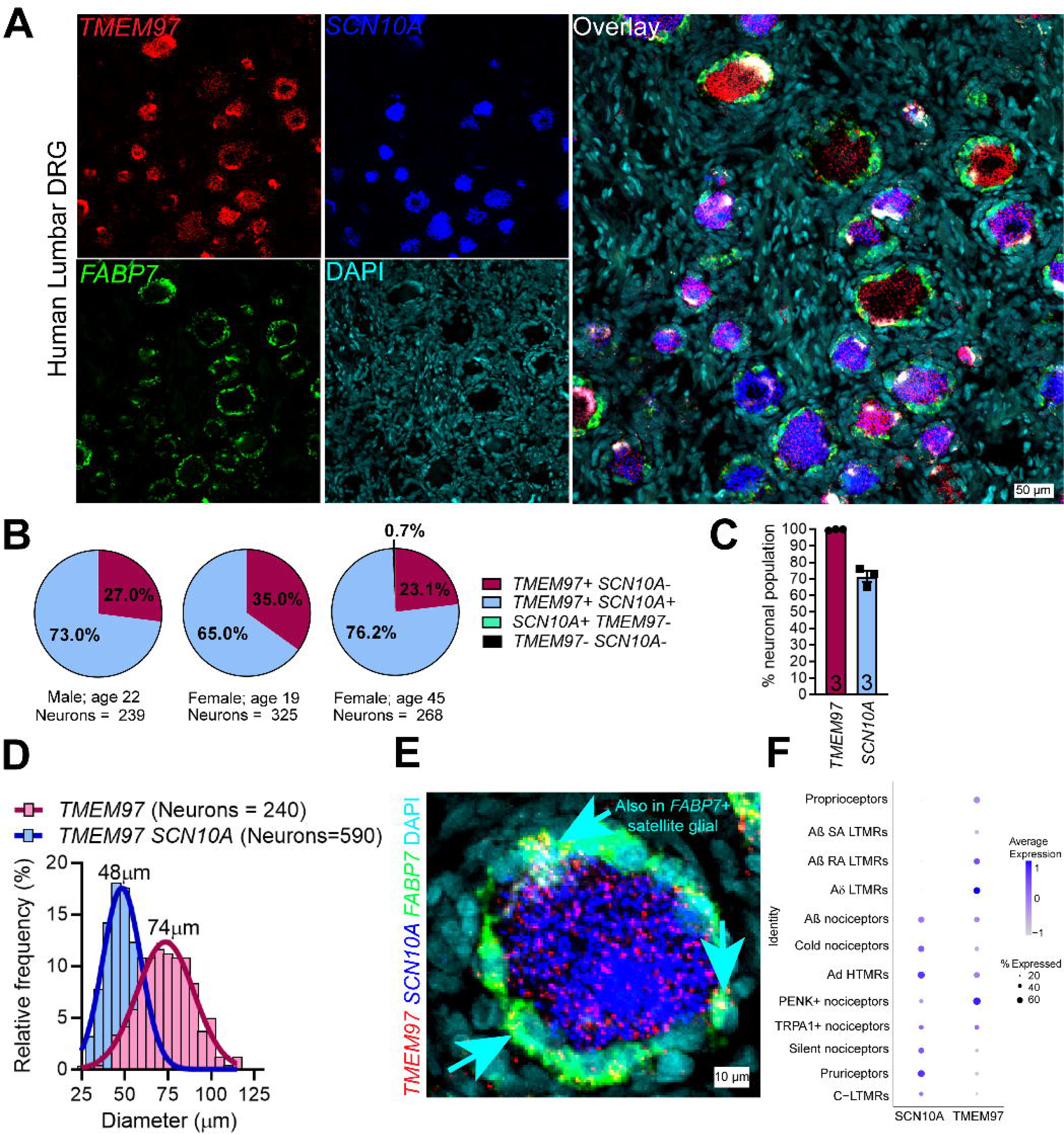
*TMEM97* gene is expressed in human dorsal root ganglia (DRG). (A) RNAScope *in situ* hybridization experiments using lumbar DRGs obtained from organ donors. (B-C) Across 3 donors (1 male and 2 females), we discovered that nearly all DRG neurons (>99%) expressed *TMEM97* and notably, all *SCN10A*-positive nociceptors expressed *TMEM97*. (D) *TMEM97*-positive neurons were distributed across all cell sizes. (E) Upon further investigation, we also identified *TMEM97* transcripts in *FABP7*-positive satellite glial cells. (F) Our previously published (36) analysis of near single-cell RNA sequencing of human DRGs showed that *TMEM97* is expressed across all neuronal cell types in the ganglia including nociceptors, low-threshold mechanoreceptors (LTMRs), and proprioceptors. *TMEM97* transcripts were notably enriched in proenkephalin (PENK)+ nociceptors and Aδ LTMRs.

### Discovery of FEM-1689 as a potent σ_2_R/TMEM97 binding ligand

Methanobenzazocines and norbenzomorphans represent two distinct chemotypes of biologically active ligands that bind selectively to σ_2_R/TMEM97 (**Supp Fig 2**) (39, 40). The first group comprises UKH-1114, which alleviates mechanical hypersensitivity in a mouse model of neuropathic pain (3) and reduces neuronal toxicity in a model of Huntington’s disease (20). Based on the positive outcomes using UKH-1114 as a treatment in models of neuropathic pain, we modified its chemical structure to identify less lipophilic analogs with improved binding profiles and physicochemical properties. We synthesized FEM-1689 (**Supp Fig 3**), which has one less methylene group than UKH-1114, and found it to be a highly selective compound with improved physicochemical properties (**Fig 2**). FEM-1689 is >100-fold more selective for σ_2_R/TMEM97 than 40 CNS proteins except for σ_1_R (10-fold) and norepinephrine transporter (NET; 24-fold) (**Supp Table 1**).

**Figure 2.**
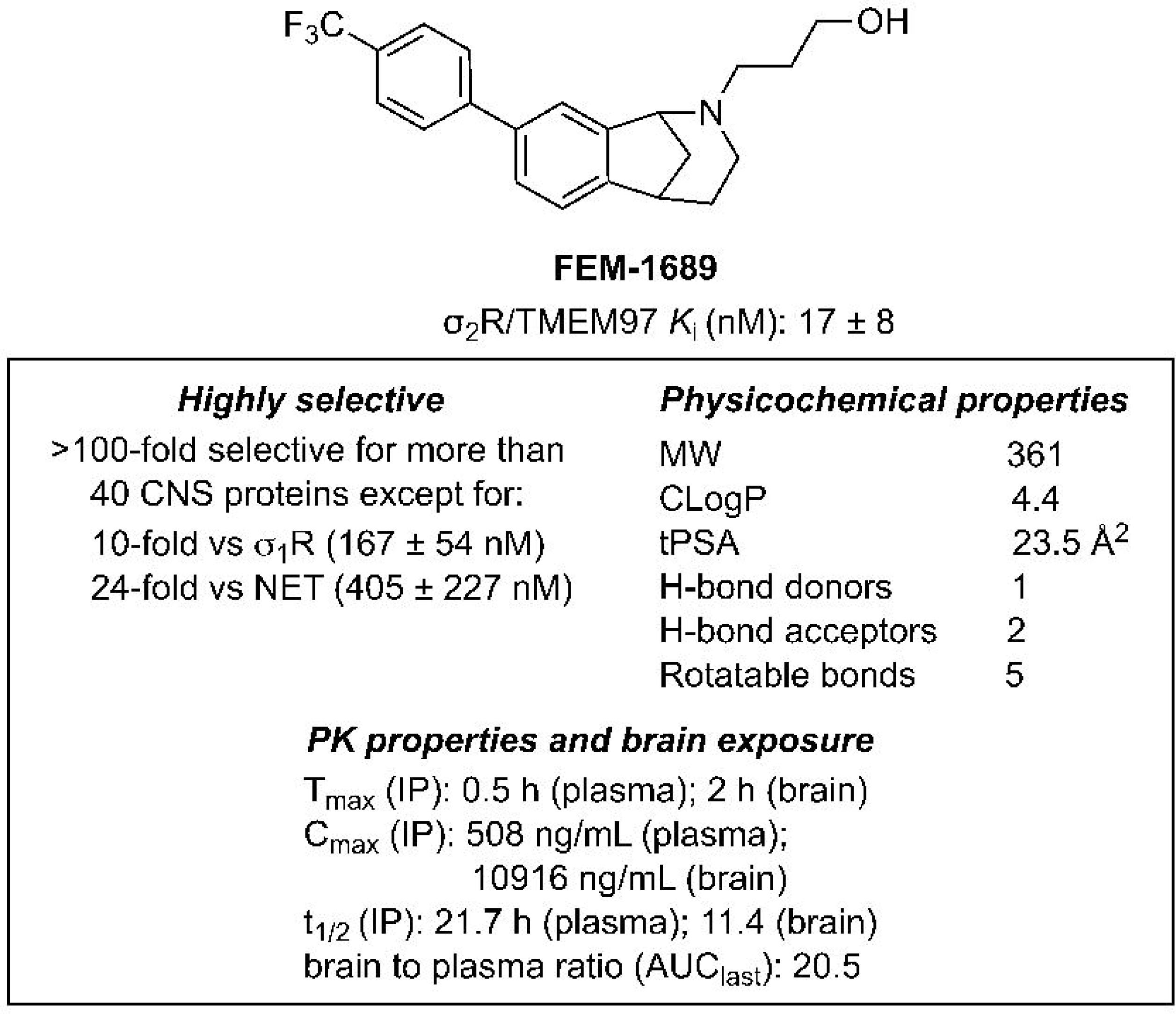
Structure, binding profiles, physicochemical properties, and pharmacokinetic parameters for FEM-1689. Values are reported as averages ± standard deviation.

### Anti-nociceptive effect of FEM-1689 in male and female mice requires an intact Tmem97 gene

An important unresolved issue is whether σ_2_R/TMEM97 ligands reduce pain hypersensitivity specifically through action on σ_2_R/TMEM97. To address this question, we used a global TMEM97-knockout (KO) mouse. Male and female TMEM97KO animals and their wild-type counterparts had similar paw mechanical sensitivity (von Frey), paw heat sensitivity (Hargreaves assay), and paw cold sensitivity (acetone) responses suggesting that the loss of TMEM97 did not alter their baseline sensation (**Supp Fig 4**). We then examined whether TMEM97KO animals develop an enhanced or blunted mechanical neuropathic pain phenotype following SNI (41) (**Fig 3A**). Male and female TMEM97KO and wild-type animals developed severe and prolonged mechanical hypersensitivity following SNI when tested at days 7, 10, and 14 after nerve injury (**Fig 3B, C**). When animals were tested 30 days following nerve injury, they were still mechanically hypersensitive. Male and female wild-type and TMEM97KO animals were then treated with a single intravenous injection of FEM-1689 dosed at 10 mg/kg, a dose determined based on previous studies with the structurally similar σ_2_R/TMEM97 ligand, UKH-1114 (3). We assessed TMEM97KO and wild-type evoked mechanical thresholds daily for 7 days and found little or no change for mechanical sensitivity in either group (**Fig 3D, E**). Animals were allowed to recover for two weeks before a second treatment with a higher 20 mg/kg dose of FEM-1689. We found that this higher dose effectively reversed mechanical hypersensitivity in both male and female wild-type mice for roughly 4 days with a single administration. Notably, FEM-1689 failed to reduce mechanical hypersensitivity in TMEM97KO mice, showing that the effects of FEM-1689 are dependent on TMEM97. The effect size measurements for FEM-1689 were quantified for males (**Fig 3F**) and females (**Fig 3G**).

**Figure 3.**
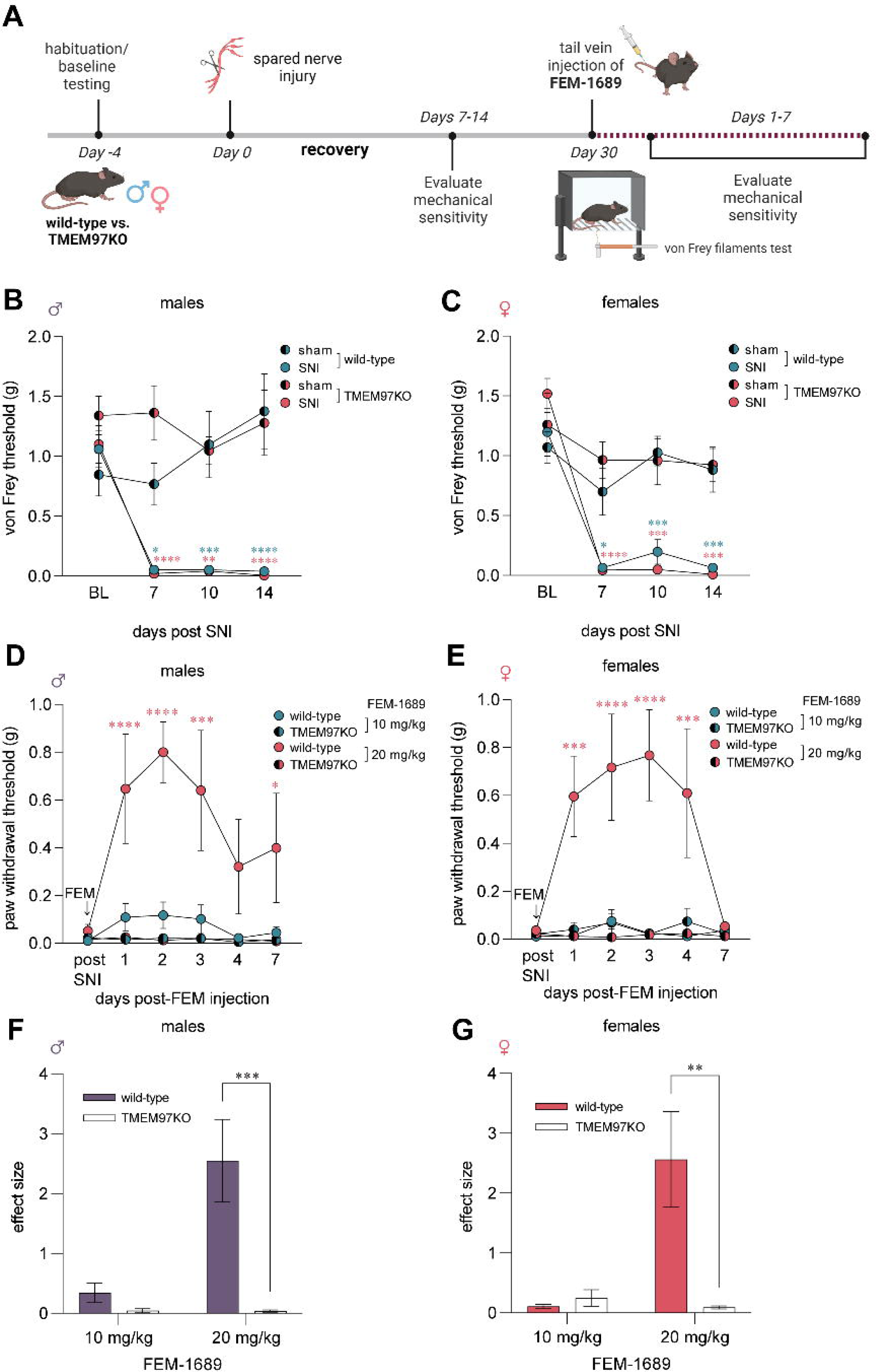
Mechanical pain hypersensitivity following spared nerve injury in wild-type and global TMEM97-knockout (KO) mice. (A) Experimental paradigm. Mechanical hypersensitivity was assessed using the von Frey filaments test at baseline prior to spared nerve injury (SNI) and for 14 days post-surgery. Mice were treated with FEM-1689 (10 mg/kg) intravenously thirty (30) days after SNI surgery and assessed for mechanical hypersensitivity for the following 7 days. Another intravenous injection of FEM-1689 (20 mg/kg) was given 2 weeks later and mechanical hypersensitivity assessed for 7 days. (B, C) Following SNI, no significant difference in mechanical hypersensitivity between wild-type and TMEM97KO littermates of both sexes was observed, suggesting that TMEM97 did not contribute to the development of neuropathic pain following nerve injury. Wild-type SNI (male n=6, female n=5), wild-type sham (male n=6, female n=5), TMEM97KO SNI (male n=5, female n=5), TMEM97KO sham (male n=5, female n=5). (D, E) A single 20 mg/kg intravenous injection of FEM-1689 reversed mechanical hypersensitivity in wild-type (male n=6, female n=5) but not TMEM97KO (male n=5, female n=5) mice. A lower dose of 10 mg/kg was not sufficient to reduce mechanical hypersensitivity in these animals. Repeated measures two-way ANOVA with Holm-Sidak’s multiple comparison test, *p<0.05, **p<0.01, ***p<0.001, ****p<0.0001. (F, G) Effect size analysis demonstrates that a higher dose of 20 mg/kg of FEM-1689 provides maximal anti-nociceptive effect. Two-way ANOVA with Sidak’s multiple comparison test, **p<0.01, ***p<0.001. Blue and red asterisks indicate wild-type and TMEM97KO groups compared to sham controls in (B, C) and wild-type vs. TMEM97KO groups in (D, E).

### FEM-1689 inhibits the ISR and promotes neurite outgrowth in a σ_2_R/TMEM97-dependent fashion

The next phase of our investigations was directed toward elucidating the mechanism by which FEM-1689 acts on sensory neurons by binding to σ_2_R/TMEM97. Prior work has suggested a link between σ_2_R/TMEM97 and cholesterol synthesis and trafficking, processes that are heavily regulated by 5’ adenosine monophosphate-activated protein kinase (AMPK) and its substrate, acetyl-CoA carboxylase (42). AMPK agonists are also known to produce anti-nociception in mouse models (22, 43–46). We treated cultured mouse DRG neurons with FEM-1689 at concentrations covering a range (10, 100, and 1000 nM), which is consistent with target binding (*K*_i_ = 17±8 nM) over the course of 16 hours, and found no change in p-ACC levels, whereas A769662 (100 µM), a known AMPK activator, increased p-ACC levels in both wild-type and TMEM97KO neurons (**Supp Fig 5A, B**). Our observation suggests that FEM-1689 does not influence the AMPK-p-ACC pathway and that a different anti-nociceptive pathway must be involved.

We then tested the hypothesis that FEM-1689 would inhibit the ISR in DRG neurons. Multiple lines of evidence support this hypothesis: 1) σ_2_R/TMEM97 and ISR transducers like protein kinase R-like ER kinase (PERK) are located on the ER membrane (40, 47); 2) a recent report on the effect of 20(*S*)-hydroxycholesterol on σ_2_R/TMEM97 implicated the ER-golgi network (12); and 3) the ISR is engaged in trauma-induced and diabetic neuropathic pain conditions in DRG neurons (25, 26). Accordingly, we cultured mouse DRG neurons from wild-type and TMEM97KO animals, treated them with FEM-1689 over 16 hours, and measured changes in the levels of p-eIF2α using immunocytochemistry (**Fig 4A, B**). Measured with immunocytochemistry (ICC), basal levels of p-eIF2α were much lower in TMEM97KO neurons than their wild-type counterparts. Treatment of wild-type neurons with ISRIB (200nM), a well-known ISR inhibitor, reduced p-eIF2α levels to the same extent as p-eIF2α levels in vehicle- and ISRIB-treated TMEM97KO neurons (**Fig 4C**). FEM-1689 reduced p-eIF2α levels, as compared to vehicle-treated cells, in wild-type mouse DRG neurons but not in DRG neurons cultured from TMEM97KO animals (**Fig 4D, E**). We found that FEM-1689 maximally inhibited the ISR at 100 nM to the same extent as ISRIB (200 nM) (**Fig 4D**) (48). We calculated the p-eIF2α IC_50_ of FEM-1689 to be 30 nM in wild-type mouse DRG neurons, which was very similar to the binding affinity of FEM-1689 to TMEM97 (*K*_i_ = 17±8 nM).

**Figure 4.**
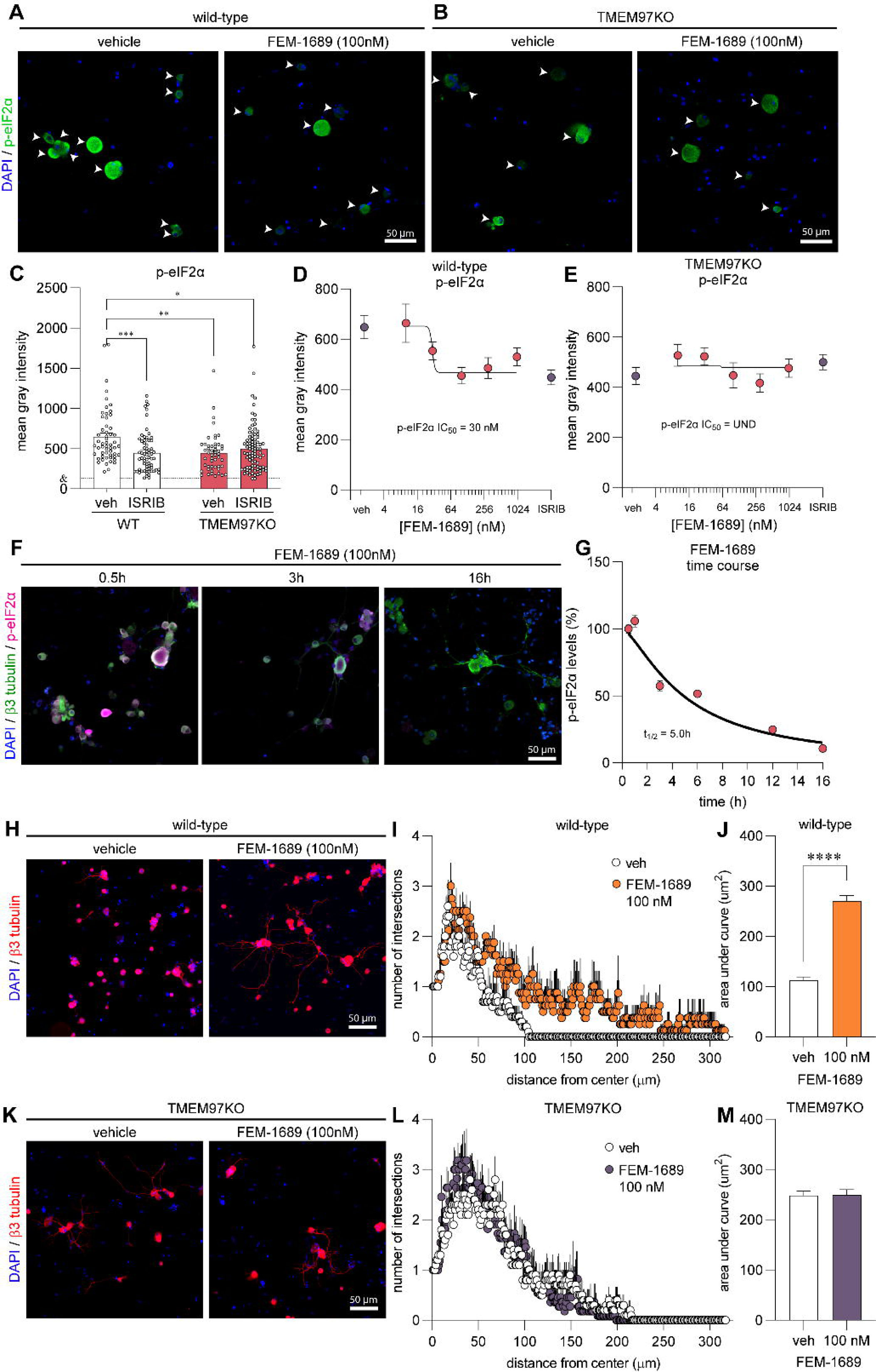
FEM-1689 reduces p-eIF2α levels and promotes neurite outgrowth *in vitro*. (A, B) Cultured mouse DRG neurons obtained from wild-type and TMEM97KO animals were treated with FEM-1689 over 16 hours. (C) Basal levels of p-eIF2α were lower in TMEM97KO neurons. ISRIB (200 nM) treatment reduced p-eIF2α levels in wild-type neurons but failed to change p-eIF2α immunoreactivity in TMEM97KO neurons. (D, E) Immunoreactivity of p-eIF2α was assessed across 5 doses (10 nM, 30 nM, 100 nM, 300 nM, and 1 µM) and a dose-response curve was generated. An IC_50_ of 30 nM was determined with maximal effect at 100 nM which was comparable to ISRIB (200 nM) treatment. TMEM97KO DRG neurons did not respond to FEM-1689 treatment. (F, G) Wild-type mouse DRG neurons were treated with FEM-1689 (100nM) at 0.5, 1, 3, 6, 12, and 16 hours. We calculated a half-life (t_1/2_) for FEM-1689 of 5.0 hours at reducing p-eIF2α levels *in vitro*. Maximal effect of FEM-1689 was observed at 16 hours of treatment. (H-M) Sholl analysis of mouse DRG neurons following 100 nM of FEM-1689 treatment showed an increase in the number and complexity of neurites in wild-type neurons but not in TMEM97KO neurons. Area under the curve of Sholl analysis was used to statistically demonstrate this effect. Immunoreactivity against β3-tubulin was used to identify neuronal cell bodies and neurites. Arrows indicate neuronal cell bodies. ***p<0.001, ****p<0.0001 two-tailed Student’s t-test. UND=undetermined. &=average mean gray intensity of the primary antibody omission control.

We further characterized the temporal dynamics of FEM-1689 in reducing p-eIF2α levels by treating wild-type mouse DRG neurons with 100 nM FEM-1689 over a time period of 0.5, 1, 3, 6, 12, and 16 hours and quantifying immunoreactivity of p-eIF2α (**Fig 4F**). Using this approach, we calculated a half-life of 5.0 hours for FEM-1689 in reducing p-eIF2α levels *in vitro*. Maximum inhibition of ISR was observed 16 hours after treatment of mouse DRG neurons with FEM-1689. This extended time period is consistent with the delayed and prolonged anti-nociceptive effects of FEM-1689. FEM-1689 did not reduce levels of BiP, a chaperone important for initiating ER stress and ISR, in either wild-type or TMEM97KO neurons (**Supp Fig 5C, D**). Sholl analysis of cultured mouse DRG neurons showed that FEM-1689 promoted neurite outgrowth in wild-type neurons but not in TMEM97KO neurons (**Fig 4F-K**). Notably, neurite outgrowth in vehicle-treated TMEM97KO neurons was more pronounced than vehicle-treated wild-type neurons – an observation that may be due to reduced ISR in TMEM97KO cells and hence, enhanced protein synthesis. These data show that σ_2_R/TMEM97 is necessary for FEM-1689 to reduce ISR and promote neurite outgrowth.

### Norbenzomorphan σ_2_R/TMEM97 ligands SAS-0132 and DKR-1677 enhance the ISR

The next question we addressed was whether other compounds that bind to σ_2_R/TMEM97 inhibit the ISR and whether ISR inhibition is specific to σ_2_R/TMEM97 modulators that promote anti-nociception. We have suggested that dissimilar biological outcomes of σ_2_R/TMEM97 modulators may arise from differences in the way individual ligands bind and interact with the protein binding pocket (40). The homologous piperazine-substituted norbenzomorphans SAS-0132 and DKR-1677 represent a chemotype that is structurally distinct from the aryl-substituted norbenzomorphan FEM-1689. Computational docking studies of SAS-0132 and FEM-1689, which were performed using the published structure of σ_2_R/TMEM97 (4), predict that these two modulators interact differently with the extended binding site of σ_2_R/TMEM97 (**Supp Fig 6**). SAS-0132, which is neuroprotective but has no anti-nociceptive effect on its own, was previously shown to inhibit the anti-nociceptive effects of UKH-1114, which is a homolog of FEM-1689 (3, 17). This observation is consistent with the hypothesis that individual σ_2_R/TMEM97 modulators may exert distinct, sometimes opposing, biological effects (3). Indeed, after treatment of wild-type mouse DRG neurons with 10, 30, 100, and 300 nM of SAS-0132 or DKR-1677 over 16 hours, we found that both compounds promote the phosphorylation of eIF2α, in contrast to the inhibitory effects of FEM-1689 (**Fig 5**). These data suggest that distinct chemotypes of TMEM97 binding compounds have differing effects on the ISR, potentially mediated by how they interact with σ_2_R/TMEM97.

**Figure 5.**
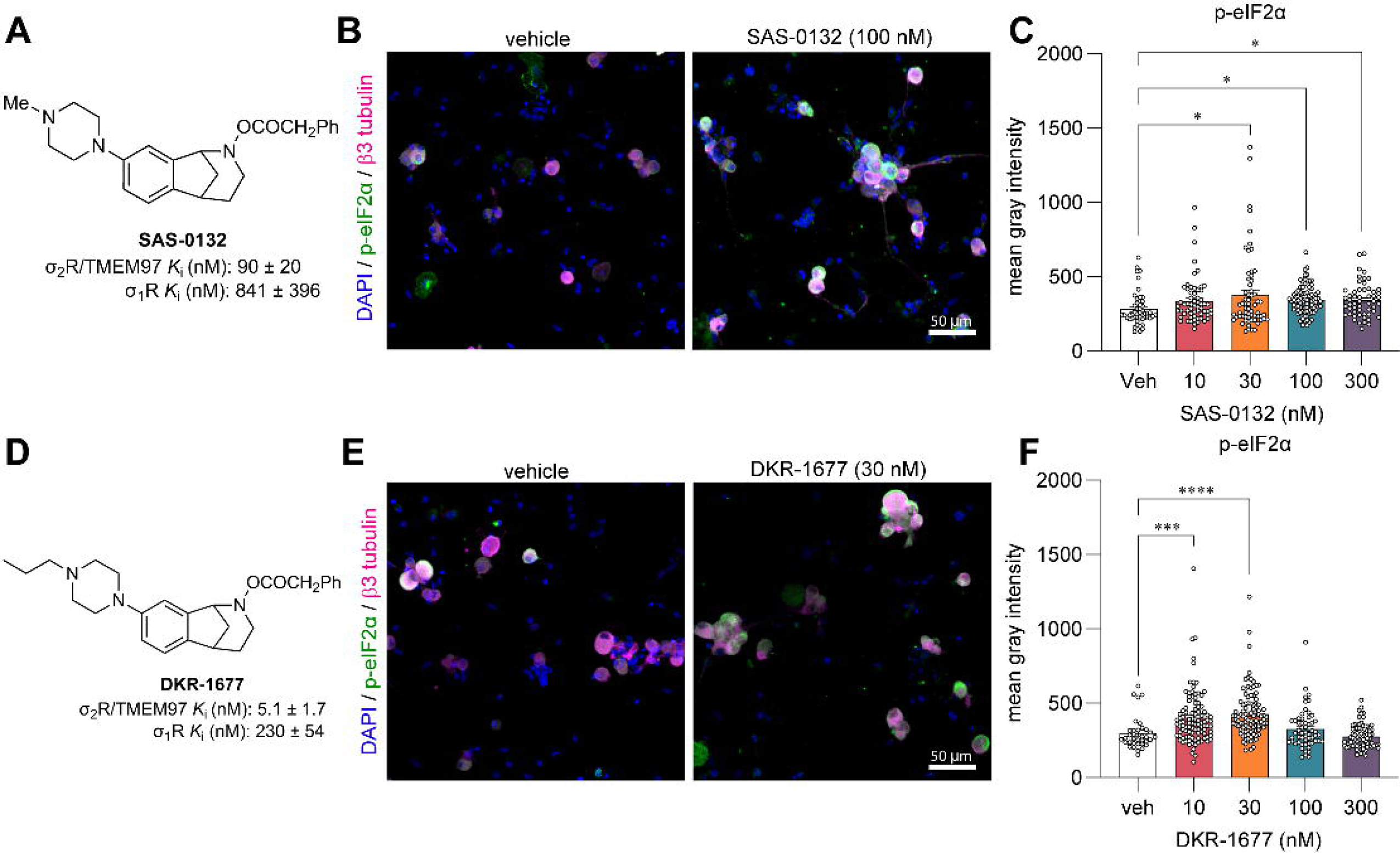
Norbenzomorphans such as SAS-0132 (A-C) and DKR-1677 (D-F) stimulate the ISR by promoting the phosphorylation of eIF2α in cultured wild-type mouse DRG neurons. Cells were treated for 16 hours with either SAS-0132 or DKR-1677 at 10, 30, 100, or 300 nM concentrations. Data is presented as fold-change compared to the fluorescence measured in the vehicle treated neurons. *p<0.05, ***p<0.001, ****p<0.0001 One-way ANOVA with Tukey’s post hoc test.

### FEM-1689 inhibits the ISR in HEK293T cells

Human embryonic kidney (HEK) 293T cells express σ_2_R/TMEM97 (The Human Protein Atlas (49)), so we queried whether FEM-1689 would inhibit the ISR in this cell line. We found a concentration-dependent reduction in p-eIF2α immunoreactivity following FEM-1689 treatment for either 2 or 16 hours using ICC and spectrophotometry (**Fig 6A, B**). We tested the effect of FEM-1689 over nine concentrations, ranging from 0.1 nM to 1000 nM and measured p-eIF2α IC_50_ of FEM-1689 in HEK cells to be 5.9 nM and 0.7 nM for 2 and 16 hour treatment conditions, respectively (**Fig 6A, B**). We found that the 2 hour FEM-1689 incubation period produced a curve with sigmoidal characteristics. We examined the effects of FEM-1689 treatment on a broader panel of ISR-related proteins using Western blotting. HEK293T cells treated with FEM-1689 over 16 hours showed a concentration-dependent reduction in phosphorylation of eIF2α, eIF2A, and p-PERK, while BiP expression remained stable (**Fig 6C-K**). Our data shows FEM-1689 influences the PERK-eIF2α arm of the ISR in HEK293T cells and that HEK293T cells can be used to develop a drug screening platform for pain therapeutics that target TMEM97.

**Figure 6.**
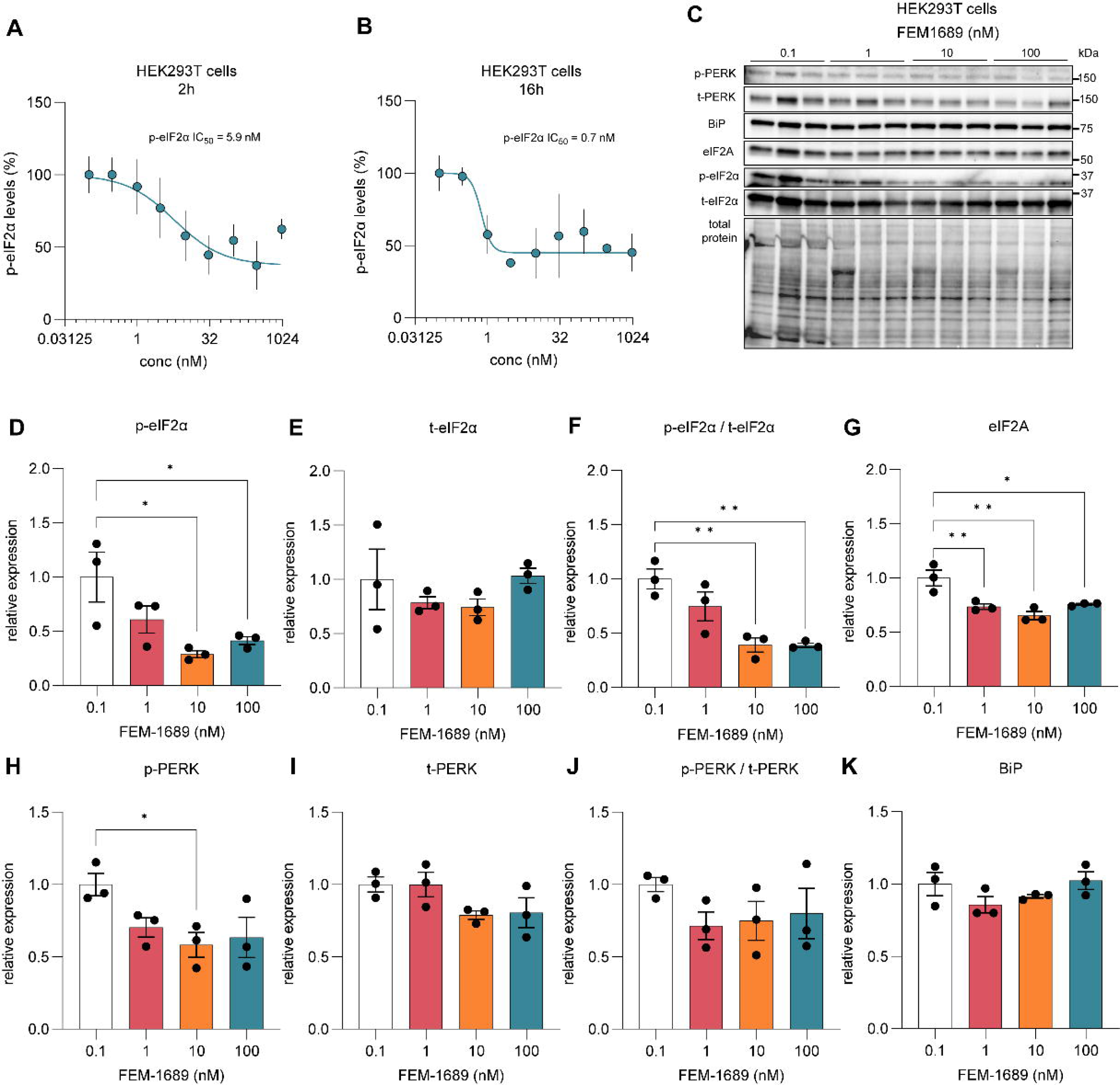
(A, B) HEK293T cells were treated for 2 and 16 hours with FEM-1689 for across a range of 9 concentrations (0.1, 0.3, 1, 3, 10, 30, 100, 300, 1000 nM) in 3-5 replicates. P-eIF2α levels were measured using ICC and spectrophotometry. IC_50_ was calculated to be 5.89 nM for the 2 hour treatment and 0.74 nM for the 16h treatment. FEM-1689 demonstrated time-dependent effect in reducing p-eIF2α levels in HEK293T cells. (C-K) In a separate experiment, HEK293T cells treated with FEM-1689 (0.1, 1, 10, 100 nM) overnight were used for Western blot analysis. Western blots showed a significant reduction in p-eIF2α, eIF2A, and p-PERK levels following FEM-1689 treatment suggesting the involvement of the PERK arm of ISR. BiP levels remained unchanged. One-way ANOVA with Tukey’s post hoc analysis *p<0.05, **p<0.01.

### FEM-1689 reverses methylglyoxal induced mechanical hypersensitivity

Methylglyoxal (MGO) is a metabolic by-product of glycolysis that is implicated in diabetic neuropathic pain and other painful neurodegenerative conditions (50–54). We have previously demonstrated that MGO induces an ISR response linked to mechanical hypersensitivity in rodents (26). We sought to determine whether the effect of FEM-1689 on reducing levels of p-eIF2α can alleviate ISR-dependent mechanical hypersensitivity caused by MGO. We treated wild-type mice with a single intraplantar injection of MGO (20 ng) to produce mechanical hypersensitivity lasting 6 days (**Fig 7A**). A group of animals were treated with either FEM-1689 (20 mg/kg, IV) or ISRIB (2.5 mg/kg, intraperitoneal, IP) 24 hours following MGO injection. FEM-1689 completely reversed MGO-induced mechanical hypersensitivity over the remaining time course of the experiment whereas the effect of ISRIB was transient (**Fig 7B**). Our data show that FEM-1689 can completely reverse ISR-dependent pain hypersensitivity caused by MGO, demonstrating the utility for developing TMEM97 modulators for diabetic neuropathic pain.

**Figure 7.**
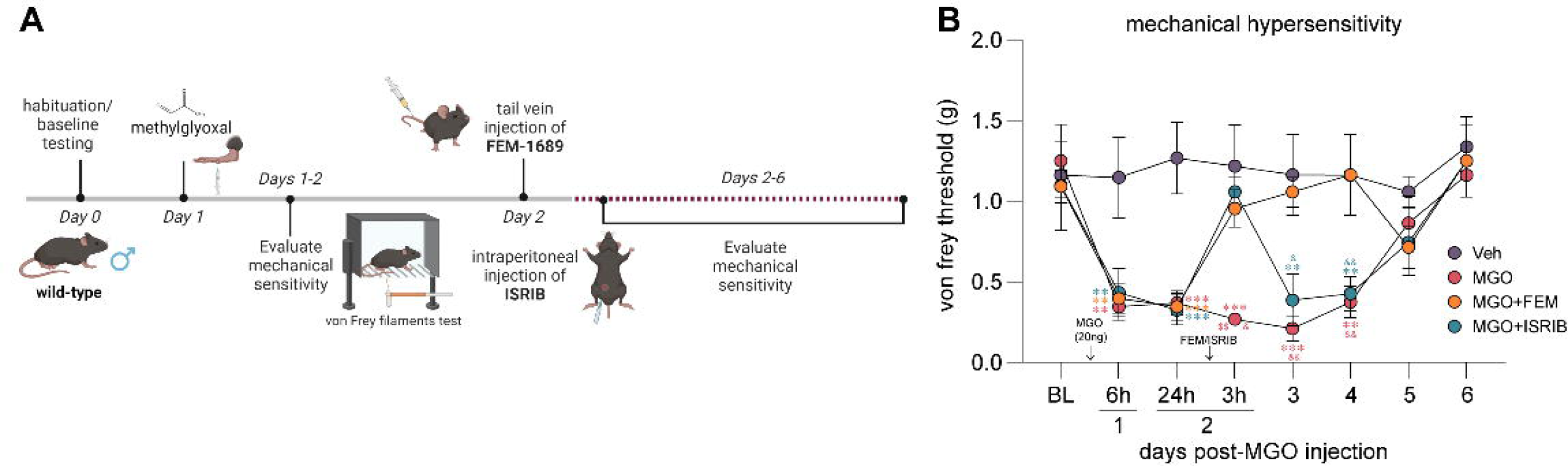
(A) Methylglyoxal (MGO, 20 ng) injection in the hind paw induces ISR-dependent mechanical pain hypersensitivity in wild-type mice over a course of 6 days. Mice were treated with FEM-1689 (20 mg/kg, IV) or ISRIB (2.5 mg/kg, IP) 24 hours following MGO administration. (B) FEM-1689 and ISRIB reversed mechanical hypersensitivity in MGO-treated mice. ISRIB’s anti-allodynic effects were observed 3 hours after drug administration and reverted to a hypersensitive state 24 hours after injection. FEM-1689 alleviated mechanical hypersensitivity for the duration of the pain state. *p<0.05, **p<0.01, ***p<0.001 repeated measures two-way ANOVA with Tukey’s post hoc test. Asterisk (*), dollar sign ($), and ampersand (&) denote post-hoc comparison with vehicle (*), MGO+ISRIB ($), and MGO+FEM (&), respectively.

### FEM-1689 reduces p-eIF2α and reverses MGO-induced ISR activity in human sensory neurons

To extend our findings with rodent DRG findings to humans, we treated cultured human DRG neurons from organ donors with FEM-1689 for 16 hours. Consistent with our mouse data, we discovered that FEM-1689 significantly reduced p-eIF2α levels in human neurons at concentrations of 10 and 100 nM (**Fig 8A, B**). We then assessed whether FEM-1689 could reverse pathological ISR activation in human DRG neurons. To induce ISR *in vitro*, we treated human neurons with MGO at 1 µM, a concentration found in the plasma of diabetic neuropathic pain patients (50). MGO treatment increased p-eIF2α levels in human DRG neurons and co-treatment with FEM-1689 (100 nM) prevented this increase (**Fig 8C, D**). These findings support the conclusion that ISR activation associated with stimuli that cause neuropathic pain in humans can be blocked by TMEM97 modulation.

**Figure 8.**
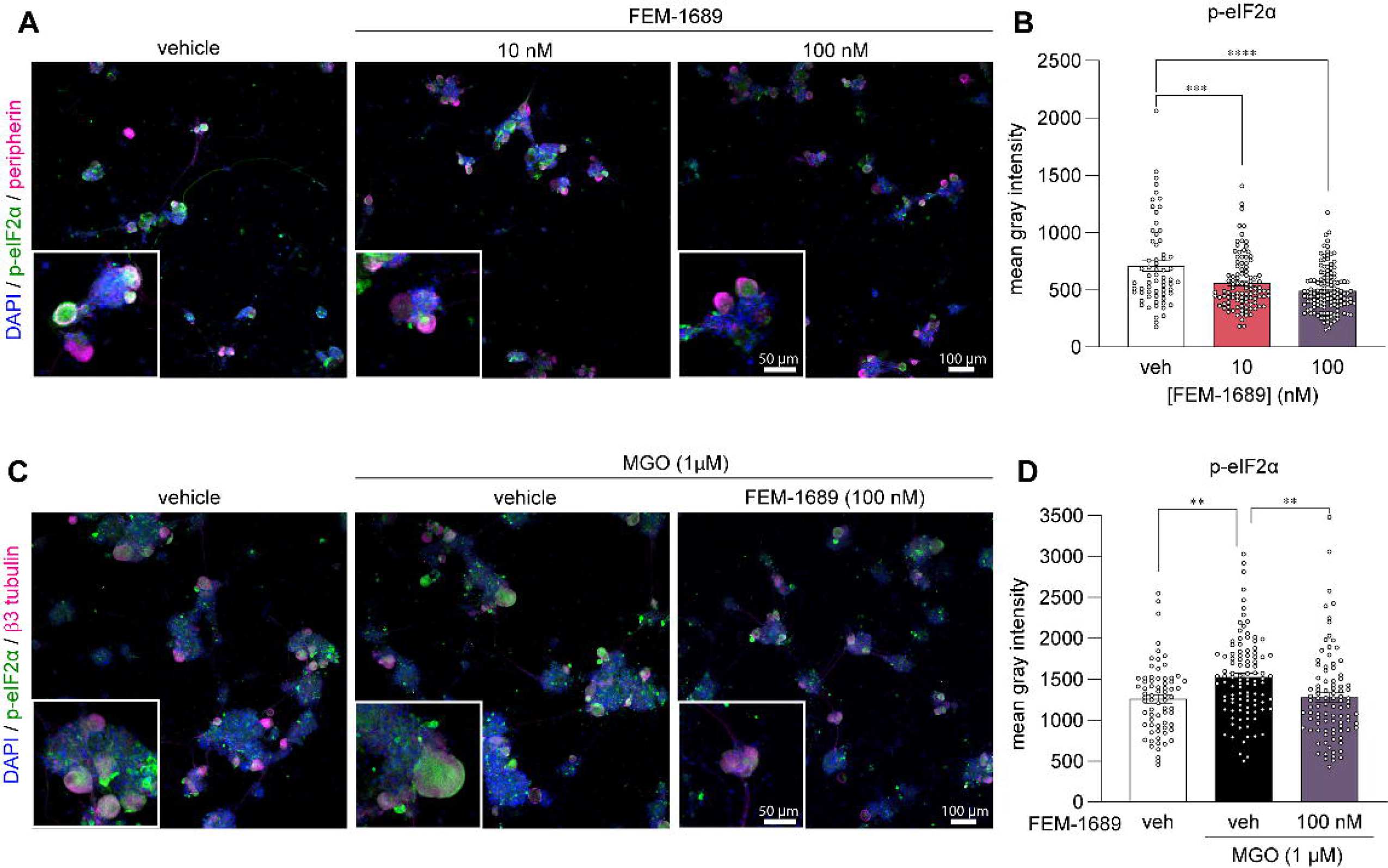
(A, B) FEM-1689 treatment (10 and 100nM) of cultured human DRG neurons significantly reduced p-eIF2α levels. (C, D) Methylglyoxal (MGO) is known to induce the ISR. Co-treatment of human neurons with MGO (1 µM) and FEM-1689 (100 nM) prevented an increase in p-eIF2α suggesting that FEM-1689 limits the effect of MGO. Peripherin and β3-tubulin were used to identify DRG neurons. One-way ANOVA followed by Tukey’s post-hoc test **p<0.01, ***p<0.001, ****p<0.0001.

## Discussion

Our experiments clearly demonstrate that the anti-nociceptive effects of a σ_2_R/TMEM97 ligand in mice of both sexes require direct modulation of σ_2_R/TMEM97, not σ_1_R or any other protein or receptor. This work also shows that modulation of σ_2_R/TMEM97 leads to the inhibition of the ISR in mouse and human DRG neurons. Because reduction of the ISR has been linked to pain relief (22, 26, 29), we posit that a plausible cellular mechanism for the anti-nociceptive effects of FEM-1689 and by extension σ_2_R/TMEM97 involves reducing the ISR. Finally, we show that human nociceptors express the *TMEM97* gene, FEM-1689 reduces eIF2α phosphorylation in cultured human DRG neurons, and that the MGO-induced ISR in human neurons can be prevented using FEM-1689. These observations suggest that targeting TMEM97 in pain patients could reduce mechanical hypersensitivity by inhibiting the ISR, a mechanism similarly observed in mice. We conclude that these findings nominate σ_2_R/TMEM97 as a *bona fide* target for developing new treatments for neuropathic pain.

The time course of action of σ_2_R/TMEM97 modulators in mouse neuropathic pain models is different from many other anti-nociceptive compounds that have a rapid onset of action. Prior to the present study, the slow onset of these anti-nociceptive effects in the mouse SNI model were independently observed by two different groups using distinct classes of σ_2_R/TMEM97 ligands (3, 4). Herein, we have shown that this effect is mediated specifically by σ_2_R/TMEM97 because the anti-nociceptive activity is completely lost in male and female TMEM97KO mice. These observations suggest that the kinetics of signaling for σ_2_R/TMEM97 may be the primary underlying reason for the delayed response. The ISR is a major mechanism that controls long-lasting changes in gene expression by influencing the translation of key injury-induced transcription factors like activating transcription factor 4 (ATF4) and C/EBP homologous protein (CHOP). ISR mediated changes involving multiple transcriptional and post-translational modifications take time to initiate and progress. It is possible that this time course is responsible for the delay in the onset of action of FEM-1689 versus the pharmacokinetics of ligand binding to σ_2_R/TMEM97. The molecular signaling pathways downstream of σ_2_R/TMEM97 modulation requires further investigation.

There is an important factor that differentiates other studies of ISR inhibitors and our findings with FEM-1689. In previous studies (26, 29), we noted that ISR inhibitors ISRIB and 4-PBA alleviate mechanical hypersensitivity over a relatively shorter duration of efficacy (hours) *in vivo* than FEM-1689 (days). The discrepancy in the duration of efficacy of ISRIB and FEM-1689 may be due to differences in their mechanism of action upstream of the ISR pathway as well as the pharmacokinetics of each compound. ISRIB allosterically binds to and promotes the catalytic activity of eIF2B, a guanine nucleotide exchange factor necessary for the recycling of non-phosphorylated eIF2 complex (55–57). In conditions of prolonged and extensive ISR activation, the effects of ISRIB are blunted (58). It is unlikely that FEM-1689 shares the same ISR inhibiting mechanism as ISRIB. So, our observations suggest that FEM-1689 may provide a distinct, more long-term way of modulating ISR than what has been observed previously with ISRIB.

Over the course of our studies to discover compounds that bind selectively to σ_2_R/TMEM97, we have discovered several chemotypes that exhibit beneficial effects in a number of animal models (40). One group comprises aryl-substituted methanobenzazocines such as UKH-1114 and JVW-1034 (**Fig 2**). UKH-1114 alleviates mechanical hypersensitivity following nerve injury (3), and it reduces neuronal toxicity induced by mutant huntingtin protein in a model of Huntington’s disease (20). The methanobenzazocine JVW-1034 not only reduces withdrawal behaviors in two rodent models of alcohol-dependence (59, 60), but it also alleviates heightened pain sensitivity that is induced by chronic alcohol exposure in mice (60). Herein we report that FEM-1689, a close analog of UKH-1114, also alleviates mechanical hypersensitivity following nerve injury and in response to MGO treatment. Another structural class of σ_2_R/TMEM97 modulators include piperazine-substituted norbenzomorphans such as SAS-0132 and DKR-1677 (**Fig 2**). For example, SAS-0132 is neuroprotective and improves cognitive performance in animal models of age-related neurodegeneration (17, 61). Notably, SAS-0132 also blocks the antinociceptive activity of UKH-1114 (3), suggesting different σ_2_R/TMEM97 binding compounds may have differing effects on nociception. DKR-1677, a homolog of SAS-0132, is protective in two different models of traumatic brain injury (TBI). It reduces axonal degeneration and provides dose-dependent enhancement of cognitive performance in the blast injury model of TBI, while it protects oligodendrocytes and cortical neurons in the controlled cortical impact model (19). DKR-1677 also protects retinal ganglion cells from ischemia/reperfusion injury (21). We demonstrate herein that SAS-0132 and DKR-1677 increase p-eIF2α expression in mouse DRG neurons, whereas FEM-1689 reduces p-eIF2α, suggesting that these compounds have distinct effects on the ISR. These observations provide the basis for developing a drug screening framework for novel pain therapeutics targeting σ_2_R/TMEM97.

Understanding how σ_2_R/TMEM97 modulators affect the ISR requires further investigation, but there are various clues for direct and indirect influence on the ISR in the literature. Firstly, σ_2_R/TMEM97 localization to the ER membrane may link it to the ISR via ER stress, particularly by influencing eIF2α phosphorylation by the kinase PERK. Indeed, we observed a reduction in p-PERK in HEK cells following FEM-1689 treatment at concentrations where p-eIF2α and eIF2A are maximally reduced (**Fig 6**), suggesting a possible link between σ_2_R/TMEM97 and the PERK pathway. Second, σ_2_R/TMEM97 is likely involved in transporting bioactive lipids, such as hydroxycholesterols (12), between cellular compartments, perhaps via a transporter activity like that of Niemann-Pick C1 (NPC1) protein (9). Excessive lipid intake and the demand for increase lipid synthesis promote lipid stress of the ER that is known to activate the PERK-eIF2α branch of the ISR and impair mitochondrial function (62–64). Finally, σ_2_R/TMEM97 regulates cellular Ca^2+^ dynamics via its influence on store-operated calcium entry (11). The ER is the largest Ca^2+^ store in the cell and is sensitive to fluctuations in Ca^2+^ levels causing ER stress and ISR induction (65). It is currently unclear whether one or multiple mechanisms linked to inhibition of the ISR are required for anti-nociception associated with FEM-1689. However, discovery of this link to ISR inhibition enables further exploration of the downstream mechanistic actions of σ_2_R/TMEM97 modulators in DRG neurons.

Neurite outgrowth can be used to assess the neuromodulatory, neuroprotective, and neuroregenerative effects of drugs (66). Our findings demonstrate that FEM-1689 promotes neurite outgrowth and complexity in a σ_2_R/TMEM97-dependent manner as measured by Sholl analysis. Neurons lacking σ_2_R/TMEM97 also display enhanced neurite outgrowth compared to their wild-type counterparts without any drug treatment. This is explained by a reduced ISR level (**Fig 5C**), and hence uninhibited protein synthesis in TMEM97KO neurons at basal levels. Axonal growth is a protein synthesis-demanding process so inhibition of the ISR in these conditions is consistent with an enhancement of protein synthesis driving this effect. Previous studies are also consistent with our findings as the ISR inhibitor ISRIB also enhances neurite outgrowth (67). Neuropathic pain models, like SNI, diabetic neuropathy, and chemotherapy-induced neuropathy are characterized by changes in axonal structure in the skin. A recent report demonstrates that aberrant reinnervation of mechanosensitive structures in the skin by nociceptors is a causative factor in late stages of neuropathic pain in mice (68). While we have not assessed skin innervation in the models we have tested, we have previously observed efficacy for σ_2_R/TMEM97 modulators in the SNI model at late time points where these skin innervation changes are observed (3). Given the effect we have seen on neurite outgrowth, it will be important to assess whether such effects can be caused *in vivo* with σ_2_R/TMEM97 ligands and whether they can lead to appropriate reinnervation of skin by nociceptors, avoiding aberrant targeting of Meissner’s corpuscles that is associated with neuropathic pain (68).

The findings reported herein provide new insights regarding the mechanistic origin of the anti-nociceptive effects that can arise from modulating σ_2_R/TMEM97 with small molecules. There remain, however, unanswered questions. First, a definitive link between ISR activation and anti-nociception *in vivo* with FEM-1689 must be established. One way to do this would be to evaluate neuropathic pain in *Eif2s1*-knockout mice. Such an experiment is complicated because loss of eIF2α, encoded by *Eif2s1*, or completely preventing the phosphorylation of eIF2α through a point mutation is lethal (69). Moreover, no specific antagonists of eIF2α have been described. The widely used classes of compounds that inhibit ISR are chemical chaperones, ISR kinase inhibitors, and eIF2B-stabilizers like ISRIB that do not directly target eIF2α and are not appropriate tools to answer this question. Secondly, it is important to acknowledge that FEM-1689 may have some polypharmacology as it appreciably binds to σ_1_ receptor (Ki = 167 ± 54 nM) and NET (Ki = 405 ± 227 nM), albeit with 10- and 24-fold less potency than TMEM97 (Ki = 17 ± 8 nM). Since the anti-nociceptive effect of FEM-1689 in TMEM97KO mice is completely abrogated, we conclude that the effect of FEM-1689 *in vivo* is mediated by TMEM97 and not σ_1_ receptor, NET, or other receptors. Moreover, σ_1_ receptor antagonists and NET inhibitors alleviate pain hypersensitivity over an acute (hours) or extensively prolonged (days to weeks) time period, respectively, neither of which fit our observations with FEM-1689 (33, 34, 65, 70–73). Nevertheless, at this stage, we cannot rule out the possibility of some polypharmacology with certainty. Thirdly, it is important to determine whether the activity of FEM-1689 is due to a peripheral or central site of action. While our pharmacokinetic experiments show that the compound readily enters the brain (**Fig 2**), we have also shown that FEM-1689 inhibits the ISR in the DRGs of mice and humans *in vitro*. Future experiments will use a nociceptor-specific TMEM97 knockout mouse under development to address this important issue.

## Methods

### Animals

TMEM97KO mice were donated by Dr. Liebl (University of Miami, Mouse Resource & Research Centers MMRRC, Tmem97^tm1(KOMP)Vlcg^, stock #050147-UCD) (21). These mice were back crossed with C57BL/6 mice obtained from Charles River. A colony of these animals were maintained at University of Texas at Dallas. Knockout and wild-type animals were obtained from the same colony. Genotyping was performed using PCR (WT: Fwd: CCAATCCTCTACACACTCCTGTT Rev: CTGGTGGCCGTCCCTATTT, Mutant: Fwd: ACTTGCTTTAAAAAACCTCCCACA. Rev: TCCTTCCCTGTAACCCATTTCTGGC). Animals were maintained on a 12-hour light-dark cycles. Adult male and female mice at least 12-week-old were used throughout the experiment. All experiments were in accordance with the National Institutes of Health guidelines and the Animal Care and Use Committee at the University of Texas at Dallas (protocol # 14-04).

### Spared nerve injury model of neuropathic pain

Spared nerve injury (SNI) was performed by exposing the sciatic nerve and transecting the peroneal and tibial branches of the nerve, leaving the sural branch intact. Cut nerves were ligated using 4-0 sutures (Kent Scientific #SUT-15-2) and the wound was closed using surgical staples. In sham animals, an incision was made to expose the nerves, but the nerves were not transected. SNI and sham surgeries were completed on the same day in a randomized order. SNI and sham data were obtained from two separate cohorts of nerve injury experiments. Animals were anesthetized with isoflurane/oxygen (50:50) mixture delivered through a nose cone. We ensured that pain reflexes were lost prior to the surgery. A 100 µL subcutaneous injection of 10% gentamicin (Sigma-Aldrich #G1272) was used to prevent infections. Animals were monitored daily and tested for mechanical sensitivity at days 7, 10, and 14 post-surgery. Behavioral testing was technically completed blinded to surgical treatment (SNI vs sham) although SNI animals often exhibit some cupping of the paw making a full blinding impossible.

### FEM-1689 intravenous administration

FEM-1689 was diluted in 100% dimethyl sulfoxide (DMSO, Fisher #67-68-5) to a concentration of 200 mM and stored at −20°C. It was further diluted in sterile 0.9% saline. The lateral tail vein was injected at 10 mg/kg or 20 mg/kg in 5µL per gram of mouse (100 µL for a 20-gram mouse) using a Hamilton syringe and 27-gauge needle. Animals were anesthetized with isoflurane/oxygen (50:50) during the injection. For behavioral experiments, all animals were injected with FEM-1689.

### Mouse pain behavior assays

Baseline testing was completed four days prior to SNI/sham surgery (**Fig 4**). Animals were habituated for at least an hour in their apparatus before being tested. Experimenters were blinded to genotype for all behavior assays. Following SNI, blinding to the injury was not possible because animals will SNI develop a “cupped” hind paw. Blinding to genotype was still maintained. Mechanical paw withdrawal thresholds in mice were assessed using the Simplified Up-Down (SUDO) method of von Frey filaments test (74). von Frey filaments were obtained from Ugo Basile and were calibrated in the lab using a weigh scale (VWR, 314AC) to 0.01 grams. Lateral surface of the paws were tested at baseline and following SNI. Cold and heat hypersensitivity details are outlined in **SI Appendix**.

### Mouse RNAscope in situ hybridization

For details regarding mouse RNAscope in situ hybridization experiments, see **SI Appendix**.

### Human samples – DRG culturing

In collaboration with the Southwest Transplant Alliance, human dorsal root ganglia (DRG) were obtained from organ donors (Supp Table 2). Once extracted, DRGs were either frozen in pulverized dry-ice for RNAScope experiments or immersed in ice-cold N-methyl-D-glutamate-supplemented artificial cerebrospinal fluid (as per (75)) until enzymatic dissociation. Human DRGs were cleaned of any fatty tissue and cut into small 1-mm chunks. These chunks were immersed in 5 ml of pre-warmed enzyme solution (2 mg/mL STEMzyme I, 4 µg/mL DNAse I obtained from Worthington Biochemical #LS004107 and #LS002139) in Hanks’ Balanced Salt Solution (HBSS) without calcium and magnesium (Gibco #14170161). Enzyme solution containing the tissue was immersed in a shaking water bath at 37 °C for 20 min followed by trituration with glass pipettes. This step was repeated 2 - 3 times until the tissue was homogenized. Afterwards, cells were passed through a 100 µm cell strainer and the flow-through containing the cells was gently layered onto 3 mL of 10% bovine serum albumin (BSA, BioPharm #71-040-002) in HBSS in a 15 ml falcon tube. This BSA gradient containing the cells was centrifuged at 900g for 5 min at room temperature. The pellet containing the cells was resuspended in BrainPhys media (STEMCell #05790) containing 2% SM1 (STEMCell #05711), 1% N2-A (STEMCell #07152), 1% Pen-Strep (ThermoFisher #15070063), and 1% GlutaMax (Gibco #35050061). Human DRG neurons were plated and grown on glass coverslips coated with poly-D-lysine (Sigma #P7405). Cultured DRG neurons were plated for 24 hours before being treated. Methylglyoxal (Sigma-Aldrich #M0252) and FEM-1689 were serially diluted in media.

### Human samples - RNAscope in situ hybridization

Fresh frozen human DRGs were embedded in optimal cutting temperature (OCT, TissueTek) and sectioned at 20-microns onto charged slides. RNAScope was performed using the multiplex version 2 kit as instructed by ACDBio and as previously described (76) with slight modifications. Slides were fixed in buffered 10% formalin and dehydrated in 50%, 75%, and 100% ethanol. A 20-second protease III treatment was used throughout the experiment. Fluorescein β, Cy3, and Cy5 Akoya dyes were used. The probes were: *TMEM97* (ACD #554471), *FABP7* (ACD #583891-C2), *SCN10A* (ACD #406291-C3). RNA quality was assessed for each tissue using a positive control cocktail (ACD #320861). A negative control probe (ACD #320871) was used to check for non-specific labeling. Images at 40X magnification were acquired on a FV3000 confocal microscope (Olympus) and analyzed using Olympus CellSens. Neuronal diameter was measured using the polyline tool and was drawn across the widest area of the neuronal soma. Lipofuscins were identified as dense bodies of autofluorescence and not analyzed. Only puncta distinct from lipofuscin were analyzed. A positive cell was deemed to have at least one mRNA punctum.

### Immunocytochemistry (ICC)

The immunocytochemistry protocol was based on a previously published protocol (77). Primary human and mouse cultured cells as well as HEK cells were fixed with 10% formalin for 10 min, and then washed three times with PBS (1X). Cells were blocked with 10% normal goat serum (NGS) (78) in PBS-triton X (0.1%). Antibodies were diluted in an antibody solution (2% NGS and 2% BSA in PBS-triton X). The antibodies used were as follows: p-eIF2α (1:500, Cell Signaling #3398), BiP (1:1000, Cell Signaling #3177), p-ACC Ser79 (1:500, Cell Signaling #3661), β3 tubulin (1:1000, Sigma #T8578), peripherin (1:1000, EnCor Biotechnology #CPCA-Peri) and DAPI (1:10,000, Cayman Chemicals #14285). Antibodies were incubated at 4 degrees Celsius overnight. The primary antibodies were washed with PBS-tween 20 (0.05%) three times, 10 min each. The secondary antibody Alexa Fluor 488 or Alexa Fluor 555 (Life Technologies, 1:1000) were dissolved in 2% NGS, 2% BSA in PBS-triton X and incubated at room temperature for 1 hour. The secondary antibody was washed with PBS-tween 20 (0.5%) thrice. DAPI was dissolved in PBS and added to the cells for 10 min. It was washed with PBS twice. Each experiment had a primary omission control where the primary antibody was replaced with only the antibody solution to discern any nonspecific binding of the secondary antibody. Coverslips were mounted onto microscope slides and imaged on a confocal. Images were analyzed using Olympus CellSens. Neurons were preferentially selected by their presence of β3 tubulin. ROIs were set to cover the cell soma and mean gray intensities of each cell was measured. Number of cells analyzed are detailed in **Supp Table 3** and **Supp Table 4.**

### HEK cells

Human embryonic kidney (HEK) 293T cells (ATCC # CRL-3216) were generously donated by the Campbell Lab (UT Dallas). Cell stocks were previously frozen at 1 million cells/ml in 90% fetal bovine serum (FBS, ThermoFisher #SH300880340) and 10% DMSO. Cells were grown in complete medium (10% FBS, 1% Pen-Strep in DMEM/F12 + GlutaMax (Gibco #10565-018) in T-75 flasks (Greiner Bio-One #658175). Cells were maintained at 37 °C with 5% CO_2_ and passaged twice before being plated onto either 96-well or 6-well plates for ICC/spectrophotometry and western blotting, respectively. Cells were counted using the TC20 automated cell counter (Bio-Rad #1450102) using a 1:1 dilution of trypan blue (Gibco #15250061).

In a 96-well plate (ThermoFisher #165305), cells were plated at a density of 20,000 cells per well and grown to roughly 65% confluency before being treated with FEM-1689 overnight. For 2 hour FEM-1689 treatment, HEK cells were plated at a density of 32,000 cells per well and treated at roughly 85% confluency. Three to five replicates were made per condition/dose (1/2 log steps ranging from 0.1 nM to 1000 nM). HEK cells were fixed with buffered 10% formalin for 10 min and washed three times with 1X PBS. ICC was performed as outlined previously. Cells were stained for p-eIF2α and DAPI. Immunofluorescence of HEK cells was quantified using the Synergy HTX Multimode Reader. Two filters were used: excitation 360/40nm emission 460/40 (DAPI), and excitation 485/20nm emission 528/20 (p-eIF2α-AlexaFluor 488). Wells were topped with 100 µL of 1X PBS prior to being read. DAPI and p-eIF2α reads were performed at bottom with a gain of 35 and 45, respectively. Area scans of 5×5 matrix was used with 497×497 micron point spacing. Negative control wells did not receive any primary antibody, but the remainder of the protocol was the same. Relative fluorescence units (RFUs) of the negative controls were subtracted from each well. P-eIF2α fluorescence was normalized to DAPI in order to control for the number of cells present in the well and then to the fluorescence of the vehicle treatment as a percentage. Values were plotted in a X-Y plot in GraphPad Prism. Concentrations were transformed to log2 values and data was fit to a non-linear regression curve (inhibitor vs response, variable slope, four parameters) constrained at top at 100, bottom at p-eIF2α levels at 300nM, and IC_50_>0. For details regarding Western blotting of cultured HEK cells, see **SI Appendix**.

### Mouse DRG cultures

Mouse DRGs were harvested from naïve wild-type and TMEM97KO mice euthanized following isoflurane anesthesia and decapitated. Mouse DRGs were processed and plated as outlined in the human DRG section above with minor differences. Lumbar DRGs of two male and two female mice were cultured together and plated across 4 well replicates of each condition. Mouse DRG neurons were grown in DMEM/F12+GlutaMax (Gibco #10565-018) plus 1% Pen-Strep and 1% N2-A. Cultured cells were plated on glass coverslips for 24 hours before being treated with FEM-1689. SAS-0132 and DKR-1677 were dissolved in 100% DMSO to a final concentration of 200mM and stored at −20 °C. FEM-1689, SAS-0132, and DKR-1677 were serially diluted in media. Untreated media was completely replaced with media containing each compound.

### Neurite outgrowth and Sholl analysis

Cultured mouse DRG neurons from wild-type and TMEM97KO animals were treated with either vehicle or FEM-1689 (100 nM). These cells were immunolabeled with β3 tubulin and DAPI according to the ICC protocol detailed above. Cells were imaged at 20X magnification on a FV3000 confocal microscope (Olympus) and Z-stacked in 1 µm intervals. Images were compiled as maximum Z-stack on ImageJ and transformed to 8-bit images. Sholl analysis was performed using the Neuroanatomy plugin in ImageJ according to the authors recommendations (79). Only solitary cells were used for analysis. The output for each cell was the number of intersections in 1-micron increments. The number of intersections was averaged for each condition across all cells. The area under the curve was calculated using GraphPad Prism.

### Synthetic procedures and characterization

For details regarding FEM-1689 synthesis, binding assays, and docking calculations, see **SI Appendix.**

### Data Analysis

All results are presented at mean ± standard error of mean (SEM) unless stated otherwise. Statistical differences between two groups were determined by two-tailed Student’s t-test. One-way and two-way ANOVAs with repeated measures were used when comparing more than two conditions. Tukey’s or Holm-Sidak’s post hoc analysis was performed. Specific statistical tests are detailed in figure legends. Statistical significance was set at p<0.05. Data was analyzed and graphed on GraphPad Prism 9.5.1. BioRender was used to build experimental schematics. Effect sizes for FEM-1689 effects on mechanical sensitivity were calculated for each animal and averaged for the group. Effect size = |(baseline-baseline) + (baseline-day1) +…+ (baseline-day7)|.

## Acknowledgements

This work was supported by Postdoctoral Fellowship Program of Natural Sciences and Engineering Research Council of Canada, by NIH grant R01 NS0655926 (TJP), and by NIH grant R61 NS127271 (BJK). The authors are grateful to the organ donors and their families for the gift of life and research provided by their organ donation. The authors thank Anna Cervantes, Geoffrey Funk, and Peter Horton of the Southwest Transplant Alliance for human tissue recovery. We are grateful to the Robert A. Welch Foundation (F-0652) for funding. We also thank the UT Austin Mass Spectrometry Facility for high resolution mass spectral data and the NIH (Grant Number 1 S10 OD021508-01) for funding the Bruker AVANCE III 500 MHz spectrometer used to characterize new compounds. We are also grateful for the Ki determinations that were generously provided by the National Institute of Mental Health’s Psychoactive Drug Screening Program, Contract # HHSN-271-2018-00023-C (NIMH PDSP). The NIMH PDSP is Directed by Professor Bryan L. Roth at the University of North Carolina at Chapel Hill and Project Officer Jamie Driscoll at NIMH, Bethesda MD, USA.

