## Supplemental Information for "Highly specific σ_2_R/TMEM97 ligand alleviates neuropathic pain and inhibits the integrated stress response"

**Supplementary Figures and Tables:**

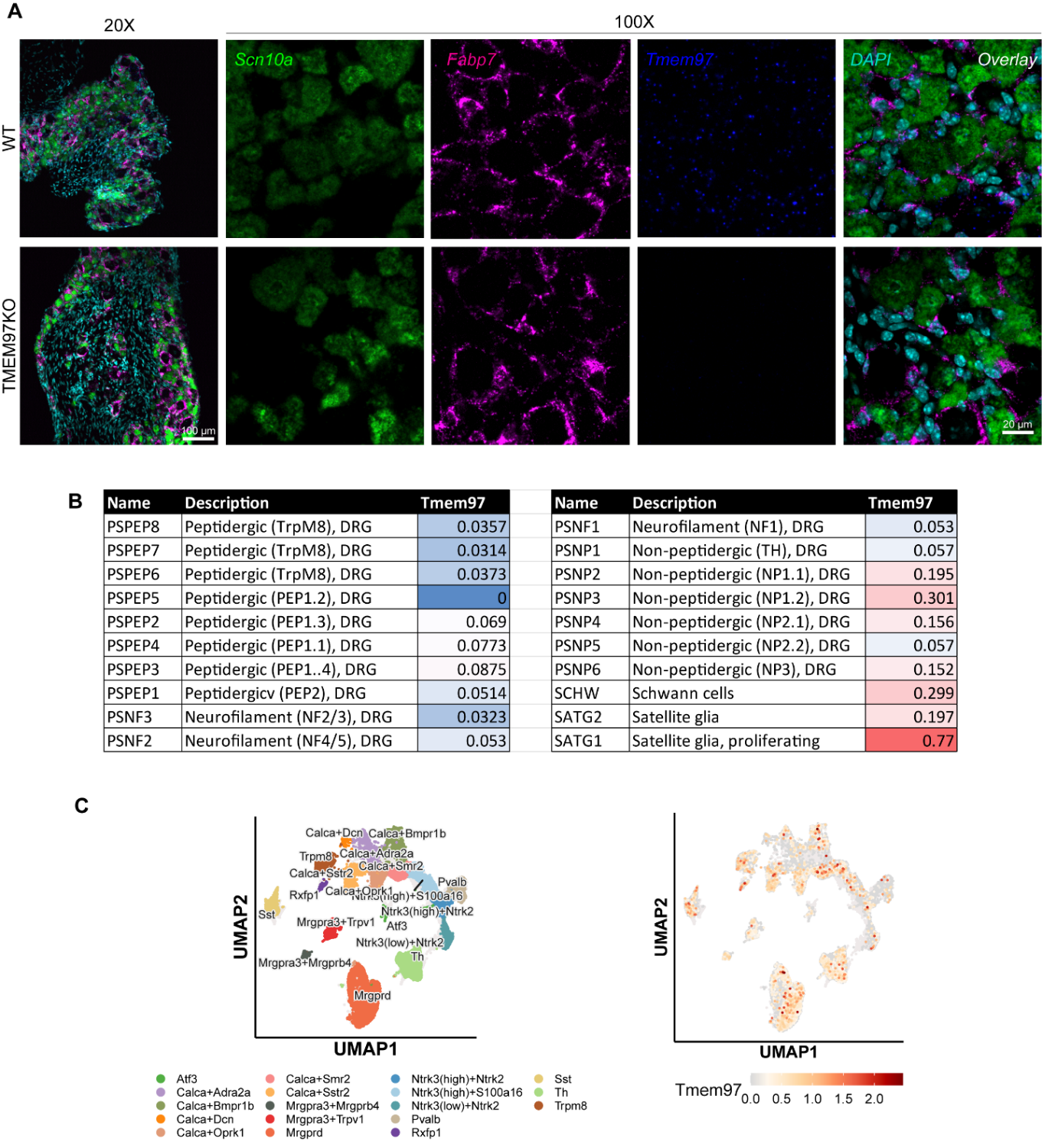

**Supp Fig 1.** RNAScope *in situ* hybridization of *Tmem97*, *Fabp7*, and *Scn10a* transcripts in mouse DRGs. (A) *Tmem97* is expressed in *Scn10a*-expressing nociceptors and *Fabp7*-expressing satellite glial cells in DRGs obtained from wild-type (WT) animals. *Tmem97* expression is lost in TMEM97-knockout (KO) DRGs. (B) Publicly available single-cell RNA sequencing dataset from mousebrain.org (1) shows that *Tmem97* is expressed across all neuronal subtypes. *Tmem97* expression is enriched in non-peptidergic neurons, Schwann cells, and satellite glial cells. (C) A recently published Harmonized Atlas of the DRG (2) integrated single-cell RNA sequencing data from humans, non-human primates, and rodents. *TMEM97* expression was found across neuronal populations in this dataset.

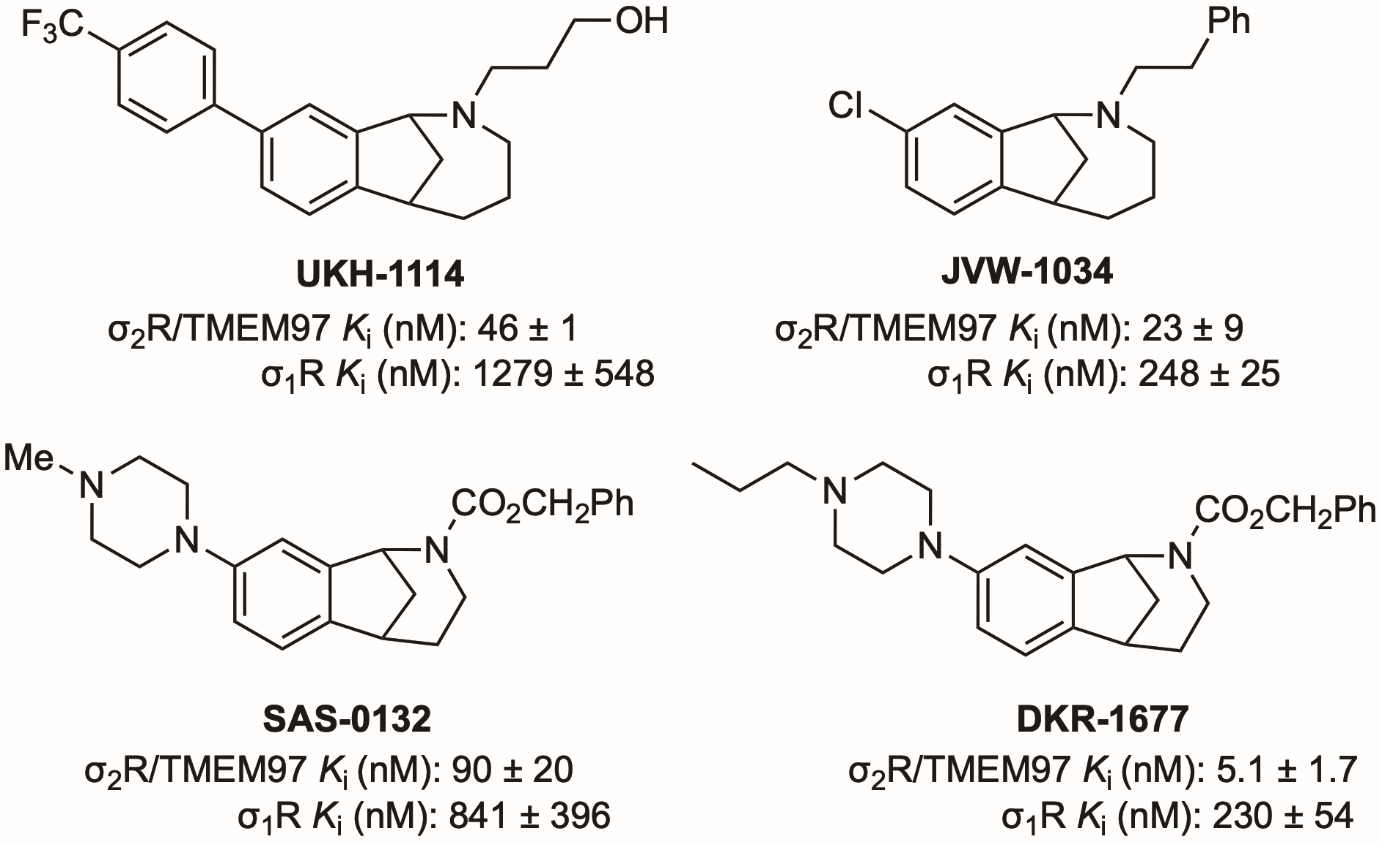

**Supp Figure 2.** Structures and binding affinities of selected biologically-active methanobenzazocines (e.g. UKH-1114 and JVW-1034) and norbenzomorphans (e.g. SAS-0132 and DKR-1677). *K*_i_ values were determined at the PDSP using σ_2_R/TMEM97 sourced from rat PC12 cells and σ_1_R sourced from guinea pig brain, and values are reported as an average ± standard deviation of two or more independent experiments (3).

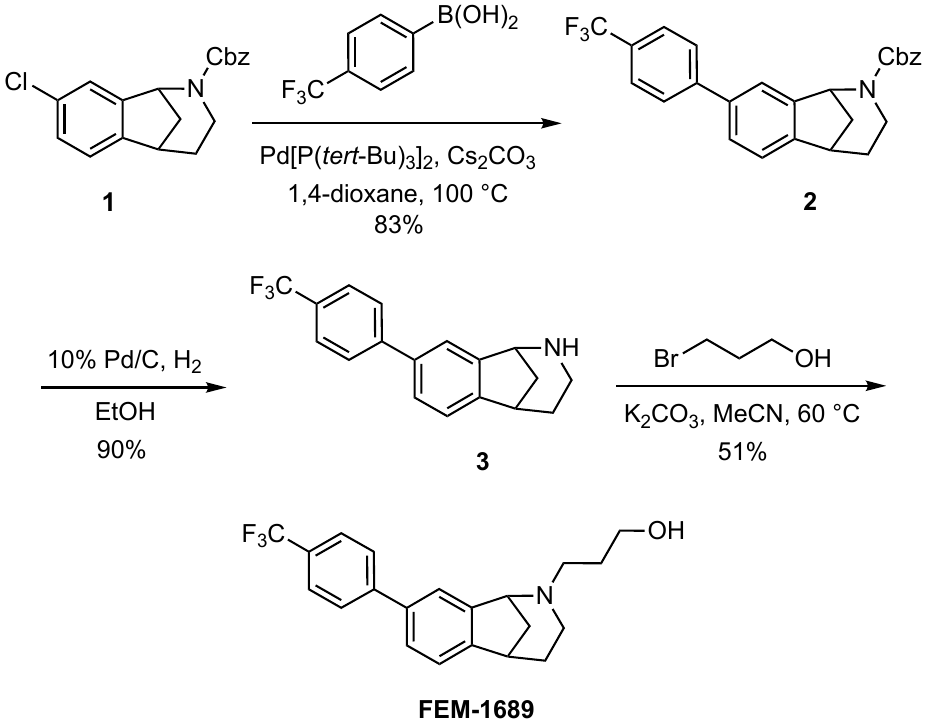

**Supp Fig 3.** Synthesis of FEM-1689.

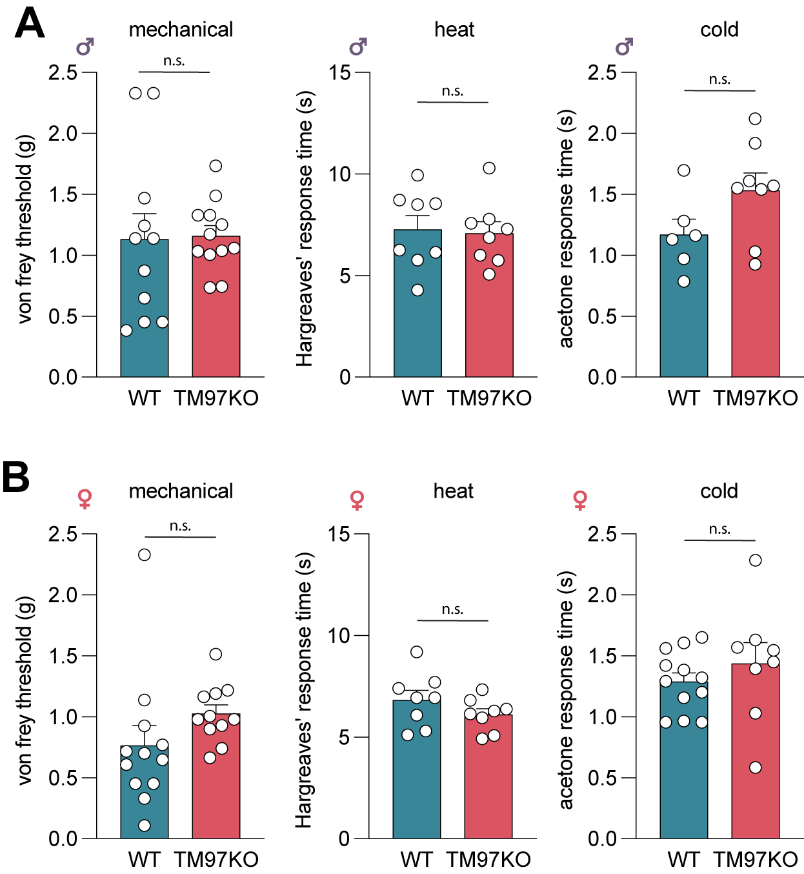

**Supp Fig 4.** Male (A) and female (B) wild-type (WT) and TMEM97KO (TM97KO) animals have similar mechanical (von Frey test), heat (Hargreaves’ test), and cold (acetone test) sensitivity under naïve, baseline conditions. A two-way student’s t-test was used to determine statistical significance: n.s. indicates “not significant”.

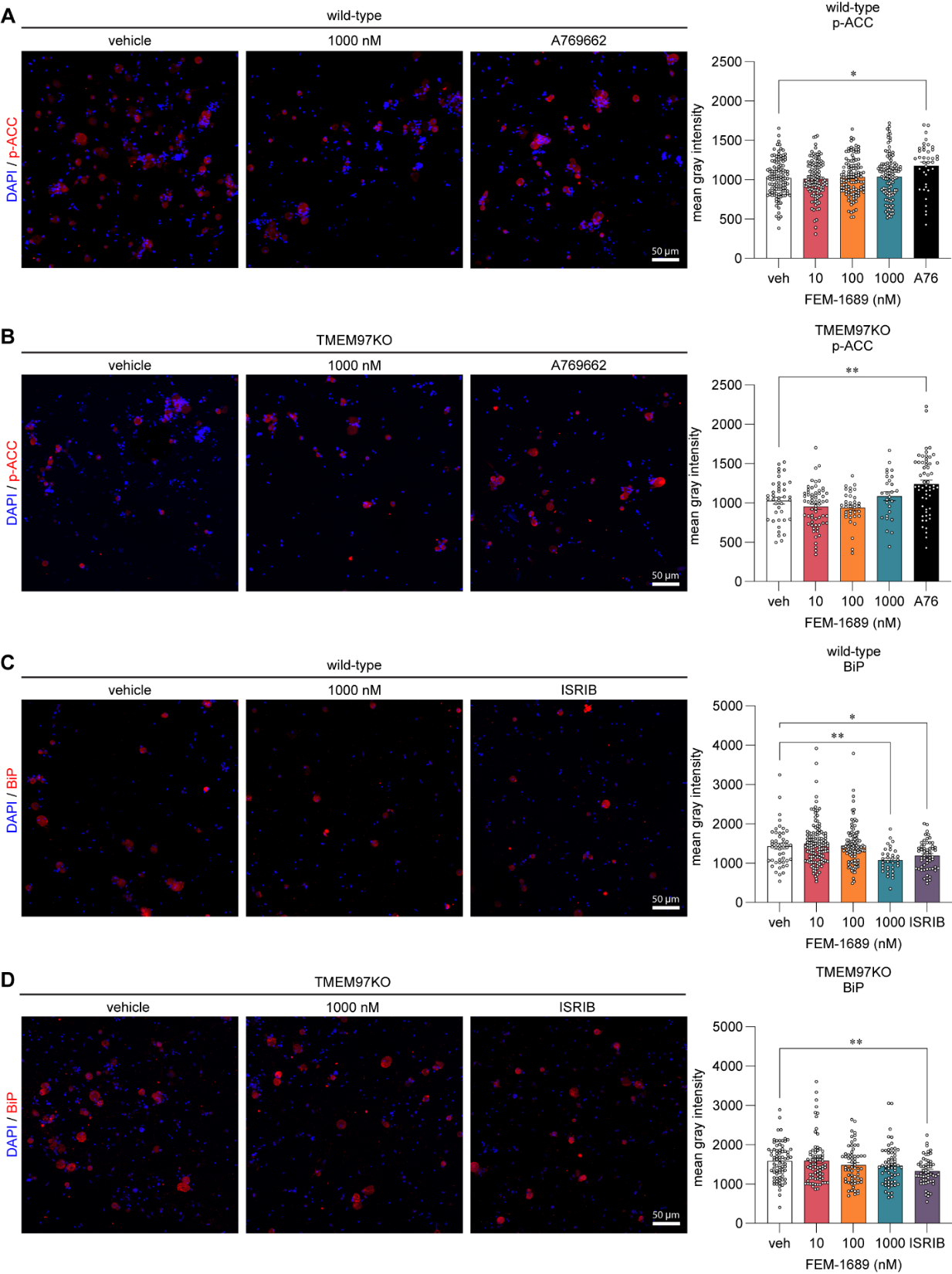

**Supp Fig 5**. Wild-type (WT) and TMEM97KO DRG neurons were treated with FEM-1689 and probed for changes in p-ACC and BiP levels using immunocytochemistry. (A, B) p-ACC levels remained unchanged in WT and TMEM97KO neurons treated with FEM-1689. A769662 (100 µM), an AMPK agonist, was able to elevate p-ACC levels in both wild-type and TMEM97KO neurons. (C, D) BiP levels in wild-type neurons were only reduced at a high concentration of 1000 nM of FEM-1689. BiP levels in TMEM97KO neurons were unaffected. ISRIB (200 nM) treatment was able to reduce BiP levels in both wild-type and TMEM97KO neurons. One-way ANOVA followed by Tukey’s post hoc test *p<0.05, **p<0.01.

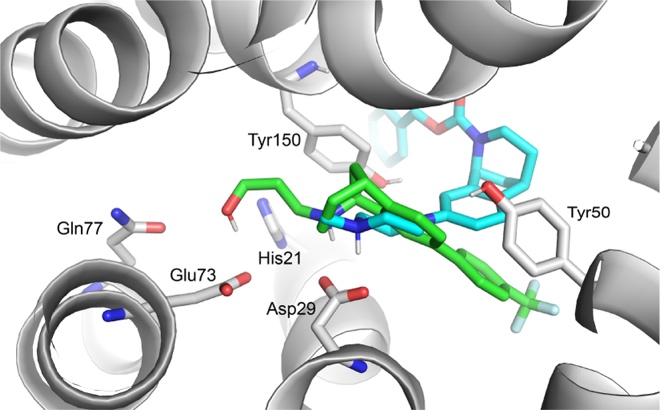

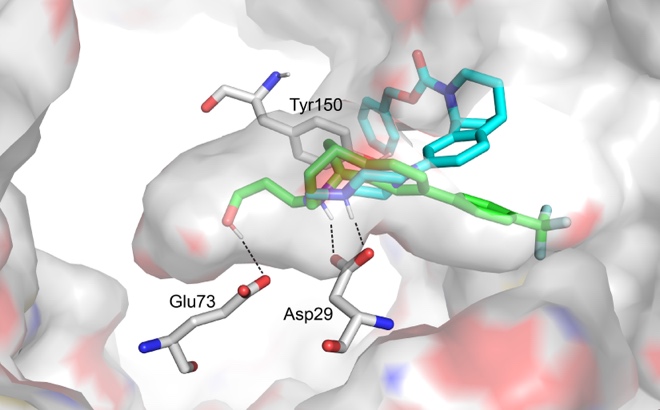

**A**

**B**

**Supp Fig 6.** Glide docking of the predicted top ranked poses of 1*S,*5*R*-enantiomers of norbenzomorphans SAS-0132 and FEM-1689 in the σ_2_R/TMEM97 binding site with selected residues shown. The σ_2_R/TMEM97 structure used for docking was the 2.71 Å structure of the bovine receptor bound to roluperidone (7M94); Chain C was used for docking studies (4). Nitrogen and oxygen atoms are colored blue and red, respectively, and the protein is represented by grey surface or cartoon and sticks for selected residues. With the exception of some polar hydrogen atoms on selected residues, hydrogen atoms on the protein are not shown, and only polar hydrogen atoms of the ligands are shown. In the binding pocket of σ_2_R/TMEM97, each ligand adopts a bound conformation in which the carboxyl group of Asp29 is positioned in close proximity to the protonated nitrogen atom of the ligand. The protonated amine in each ligand is also predicted to engage in a conserved cation-π interaction with Tyr150. Despite the similar electrostatic interactions between the protonated amino groups with Asp29 and Tyr150, other interactions between SAS-0132 and FEM-1689 within the internal binding pocket and with the external surface of σ_2_R/TMEM97 are predicted to be significantly different. The bound poses of SAS-0132 and its homolog DKR-1677 are virtually identical (data not shown). (A) Overlay of (1*S*,5*R*)-SAS-0132 (cyan) and (1*S*,5*R*)-FEM-1689 (green) docked with σ_2_R/TMEM97 showing the protein surface (grey, red, blue), the binding pocket, the protonated nitrogen atoms of the ligands, and the side chains of protein residues Asp29, Glu73 and Tyr150. Interactions between Asp29 with the protonated amino groups of ligands are shown by dashed black lines. A weaker interaction between the hydroxyl group on the side chain of FEM-1689 with Glu73 is also shown by a dashed black line. (B) Overlay of (1*S*,5*R*)-SAS-0132 (cyan) and (1*S*,5*R*)-FEM-1689 (green) docked with σ_2_R/TMEM97 showing the protein backbone (grey cartoon), the protonated nitrogen atoms of the ligands, and the side chains of protein residues His21, Asp29, Tyr50, Glu73, Gln77, and Tyr150, which are key residues in the binding pocket.

**Supp Table 1. Binding profile of FEM-1689 at non-sigma receptor sites.** Competitive binding assays were completed by the Psychoactive Drug Screening Program (PDSP) at the University of North Carolina at Chapel Hill. Percent inhibition of radioligand at targets is determined for FEM-1689. Ki’s are determined with secondary assay for any hits with greater than 50% binding in the primary assay. hERG binding is included as a safety/nuisance target.

| Target | *K_i_* (nM) | Target | *K_i_* (nM) |
| --- | --- | --- | --- |
| 5HT_1A_ | * | Beta3 | * |
| 5HT_1B_ | * | BZP Rat Brain | 6595 |
| 5HT_1D_ | * | D_1_ | * |
| 5HT_1e_ | * | D_2_ | * |
| 5HT_2A_ | * | D_3_ | * |
| 5HT_2B_ | * | D_4_ | * |
| 5HT_2C_ | * | D_5_ | * |
| 5HT_3_ | * | DOR | * |
| 5HT_5a_ | * | GabaA | * |
| 5HT_6_ | >10,000 | H_1_ | * |
| 5HT_7_ | * | H_3_ | * |
| Alpha_1a_ | 1994 | H4 | 5512 |
| Alpha_1b_ | * | KOR | 7091 ± 2395^a^ |
| Alpha_1d_ | * | M_1_ | * |
| Alpha_2a_ | * | M_2_ | * |
| Alpha_2b_ | * | M_3_ | * |
| Alpha_2c_ | * | M_4_ | * |
| Beta1 | * | M_5_ | * |
| Beta2 | * | MOR | * |
| PBR | * | NET | 405 ± 227^b^ |
| SERT | * | hERG | 839 |

* < 50% inhibition of radioligand binding at 10 µM test article.

^a^ Average of two *K*_i_ determinations; ^b^ Average of four *K*_i_ determinations; all other *K*_i_ values

are from a single determination

**Supp Table 2**. Demographics of organ donors and their cause of death.

| **ID** | **Sex** | **Age** | **Cause of Death** | **Experiment** |
| --- | --- | --- | --- | --- |
| AIGD395 | F | 45 | CVA/Stroke | RNAScope (Fig 1 A-E) |
| AJEK100 | M | 22 | Overdose/CVD | RNAScope (Fig 1 A-E) |
| AJHX028 | F | 19 | Anoxia/Overdose | RNAScope (Fig 1 A-E) |
| AJAM174 | F | 25 | Head Trauma/MVA | Cultured DRG neurons (Fig 6 A-B) |
| AJCI494 | F | 40 | Anoxia/CVA | Cultured DRG neurons (Fig 6 A-D) |
| AJDS185 | F | 16 | Head Trauma | Cultured DRG neurons (Fig 6 C-D) |

*MVA: motor vehicle accident, CVA: cerebrovascular accident, CVD: cardiovascular disease

**Supp Table 3**. Number of neurons analyzed for *in vitro* mouse DRG studies

| **Figure** | **Genotype** | **Condition** | **N** |
| --- | --- | --- | --- |
| Fig 4D | Wild-type | Vehicle | 55 |
|  |  | 10 nM | 22 |
|  |  | 30 nM | 47 |
|  |  | 100 nM | 36 |
|  |  | 300 nM | 24 |
|  |  | 1000 nM | 21 |
|  |  | ISRIB | 66 |
| Fig 4E | TMEM97KO | Vehicle | 50 |
|  |  | 10 nM | 33 |
|  |  | 30 nM | 65 |
|  |  | 100 nM | 34 |
|  |  | 300 nM | 39 |
|  |  | 1000 nM | 36 |
|  |  | ISRIB | 89 |
| Fig 4F-G | Wild-type | 0.5h | 197 |
|  |  | 1h | 145 |
|  |  | 3h | 106 |
|  |  | 6h | 103 |
|  |  | 12h | 140 |
|  |  | 16h | 130 |
| Fig 4H-J | Wild-type | Vehicle | 10 |
|  |  | 100 nM | 8 |
| Fig 4K-M | TMEM97KO | Vehicle | 10 |
|  |  | 1000 nM | 12 |
| Fig 5A-C | Wild-type | Vehicle | 82 |
|  |  | 10 nM | 61 |
|  |  | 30 nM | 63 |
|  |  | 100 nM | 86 |
|  |  | 300 nM | 54 |
| Fig 5D-F | Wild-type | Vehicle | 83 |
|  |  | 10 nM | 101 |
|  |  | 30 nM | 102 |
|  |  | 100 nM | 57 |
|  |  | 300 nM | 80 |

**Supp Table 4**. Number of neurons analyzed for in vitro human DRG studies

| **Figure** | **Condition** | **N** |
| --- | --- | --- |
| Fig 8A, B | Vehicle | 65 |
|  | 10 nM | 103 |
|  | 100 nM | 133 |
| Fig 8C, D | Vehicle | 71 |
|  | Vehicle + MGO | 103 |
|  | 100 nM + MGO | 95 |

**Supplementary Methods:**

*Mouse pain behavior assay – Cold and heat sensitivity*

Cold sensitivity was assessed using drops of acetone applied to the plantar surface of each paw. Pain-like behaviors (i.e. vigorous shaking and licking) were timed for up to 45 seconds. Each paw was tested three times and both paws were averaged.

Heat latency to respond was measured using a Hargreaves device (5) (IITC Model 400, Life Science Inc.; Harvard Apparatus, CA) with the heated glass set at 29 °C, 40% active laser power, and a 20-second cutoff. Each paw was tested three times and both paws were averaged. Light was targeted to the lateral surface of the paws.

*Mouse RNAscope in situ hybridization*

A dorsal root ganglion (DRG) tissue (bilateral L1-L5) was collected from 10-week-old male C57BL6/J wild-type (WT) and littermate-matched TMEM97KO mice. The tissue was embedded in optimal cutting temperature (OCT, TissueTek) and was immediately flash-frozen in dry ice. The DRG were sectioned at 20 microns on a cryostat, and directly mounted onto Super Frost Plus charged slides. Slides were dried at –20 °C for 2 hours to increase tissue adherence and stored at –80 °C until used for RNAscope in situ hybridization.

RNAscope fluorescent in situ hybridization multiplex assay v2 was used as instructed by Advanced Cell Diagnostics (ACDBio Inc). Probes for *Tmem97* (ACD #527591), *Scn10a* [a marker for Nav1.8-expressing nociceptors] (ACD #426011), and *Fabp7* [a marker for satellite glial cells] (ACD #414651) were used to validate the deletion of *Tmem97* expression in TMEM97KO DRGs, and localization of *Tmem97* expression in WT DRGs. Every cohort of slides included at least one slide for negative control with no target probe (ACD #320871) and one positive control slide with three positive control target probe cocktails (ACD #320861) (62) for tissue quality check. Slides were removed from –80 °C and immediately immersed in pre-chilled (4 °C) 10% neutral buffer formalin for 15 min. The slides were rinsed twice in 1X phosphate-buffered saline (PBS, pH 7.4) and dehydrated in a series of different ethanol concentrations of 50%, 70%, and 100% (twice) for 5 min each at room temperature. The hydrophobic barrier was drawn around the section using a hydrophobic pen (ImmEdge PAP pen, Vector Labs) after briefly air drying. Slides were incubated with 1:2 diluted hydrogen peroxide in distilled water for 10 min at room temperature and washed twice in distilled water. The protease IV was applied to each section and incubated for 5 min at room temperature. Slides were washed twice in 1X PBS and RNA scope was performed immediately. A mixture of probes - *Tmem97, Scn10a, and Fabp7* – was hybridized for 2 hours at 40 °C in a humidity control tray inside a HybEZ oven. Signals for all three channels were amplified using a series of AMPs, and a TSA-based fluorescent label was developed for each channel using Opal 520, 570, and 690 dyes (Akoya Bioscience) for each channel. Slides were cover-slipped with Vectashield anti-fade mounting medium with DAPI. Images were captured using an Olympus FV 3000 confocal microscope at 100X magnification and each image used the same acquisition parameters for WT, TMEM97KO, and negative control slides. Imaging was not completed blinded to genotype.

*Western blotting*

HEK cells were plated at 60,000 cells per well onto six-well plates (ThermoFisher # 351146) for 24 hours before being treated with FEM-1689 overnight. Protein extractions and western blotting was performed as previously published (6). In brief, media from the cells was removed and cells were washed once with 1X PBS. Cells were lysed in the well with radioimmunoprecipitation assay (RIPA) buffer consisting of 25 mM Tris, 150 mM NaCl, 0.1% SDS, 0.5% Na deoxycholate, 1% Triton X-100 with protease and phosphatase inhibitor cocktails (Sigma Aldrich P8340, P5726, P0044 – 1:100 each). Cells were centrifuged at 4 °C at 14,000 rpm for 10 min. The supernatant was used for downstream analysis. Protein was quantified using the Pierce BCA Protein Assay kit (Thermo Scientific, #23225) according to the manufacturer’s instructions.

Protein samples were denatured using 4X Laemmli Sample buffer (BioRad #1610747) and heated at 95 °C for 5 min. 20 µg of protein samples were loaded onto 4-20% Stain-Free Criterion TGX gels (BioRad #5678094). Gels were run in Tris/Glycine/SDS running buffer (BioRad #1610772) at 120V for roughly 1.5 hours. Stain-Free gels were activated for 5 min prior to the transfer step. Protein was transferred for 10 min onto low fluorescence polyvinylidene difluoride (PVDF) using a TransBlot Transfer kit (BioRad #1704275) and a Transblot Turbo transfer system (BioRad). Once transferred, total protein was imaged immediately using a ChemiDoc MP system (BioRad). Blots were allowed to dry for 5 min and were blocked with 5% non-fat milk in Tris buffered saline (TBS)-Tween 20 (0.05%). Antibodies were diluted in 1% non-fat milk in TBS-Tween 20. Primary antibodies used in this study were: p-eIF2α (1:1000, Cell Signaling #3398), t-eIF2α (1:1000, Cell Signaling #9722), eIF2A (1:2500, Abcam #ab169528), p-PERK (1:500, Cell Signaling #3192), t-PERK (1:1000, Cell Signaling #3179), BiP (1:1000, Cell Signaling #3177). Goat anti-Rabbit IgG horseradish peroxidase (HRP) (H+L) (1:10,000) was used as a secondary antibody. Stained blots were imaged using enhanced chemiluminescence (ECL) on a ChemiDoc MP system.

*Synthetic procedures and characterization*

*Binding assays.* Sigma receptor binding assays for FEM-1686, which was determined to be >95% pure (LC-MS), were performed by the National Institutes of Mental Health Psychoactive Drug Screening Program (NIMH PDSP) at Chapel Hill, North Carolina (3). σ_1_R and σ_2_R/TMEM97 were sourced from HEK293T cells transfected with human σ_1_R and σ_2_R/TMEM97. σ_1_R binding affinity (*K_i_*) was determined through competition binding assays with [^3^H]-(+)-pentazocine, whereas σ_2_R/TMEM97 binding affinity (*K_i_*) was determined through competition binding assays using the radioligand [^3^H]-ditolylguanidine in the presence of (+)-pentazocine to block σ_1_R binding sites. *K_i_* values are calculated from an average of two or more independent experiments. Detailed experimental protocols are available on the NIMH PDSP website at https://pdspdb.unc.edu/pdspWeb.

*General.* Commercial reagents and solvents were used without purification unless stated, but acetonitrile (MeCN) was dried by filtration through two columns of activated molecular sieves. Glassware was dried overnight in an oven at 120 °C or flame dried under vacuum for a minimum of 5 min. All air- or moisture-sensitive reactions were performed under an atmosphere of argon or nitrogen. Reaction temperatures refer to the temperature of the heating or cooling bath. Volatile solvents were removed under reduced pressure using a Büchi rotary evaporator at 25–30 °C. Air- or moisture-sensitive reagents and all solvents were transferred using plastic syringes and steel needles using standard techniques. Proton nuclear magnetic resonance (^1^H NMR) and carbon nuclear magnetic resonance (^13^C NMR) spectra were recorded at the indicated field strength in CDCl_3_. Chemical shifts are reported in parts per million (δ) and are referenced to the deuterated solvent. Coupling constants (*J*) are reported in Hertz (7), and the splitting abbreviations used are: s, singlet; d, doublet; t, triplet; q, quartet; dt, doublet of triplets; ddd, doublet of doublets of doublets; m, multiplet; br s, broad singlet;. Capillary melting points are uncorrected. Accurate mass measurements were determined using an LC-MS system comprised of an Agilent 1260 series HPLC and an Agilent 6530 single quadrupole time-of-flight mass spectrometer. Purities of all compounds submitted for testing at PDSP were determined by LC-MS from the areas under the curves (AUC) at 214 and 254 nm. Column chromatography was performed using glass columns and “medium pressure” silica gel (Silicycle, 230-400 mesh).

*Benzyl 8-(4-(trifluoromethyl)phenyl)-1,3,4,5-tetrahydro-2H-1,5-methanobenzo[c]azepine-2-carboxylate (****2****).* A solution of aryl chloride **1** (8) (327 mg, 1.0 2.0 mmol), 4-trifluoromethylphenylboronic acid (379 mg, 2.0 mmol), Cs_2_CO_3_ (650 mg, 2.0 mmol), palladium(bis)(*t*-butyl)_3_ phosphine (25.5 mg, 0.05 mmol) in degassed 1,4-dioxane (4 mL) was stirred for 21 h at 100 °C. The reaction was cooled to room temperature and poured into water (5 mL). The mixture was extracted with CH_2_Cl_2_ (3 × 15 mL), and the combined organic layers were dried (MgSO_4_) and concentrated under reduced pressure. The crude product was purified via flash column chromatography (SiO_2_), eluting with hexane/EtOAc (50:1 to 20:1 to 13:1 v/v) to afford 367.2 mg (83%) of **2** as a pale yellow oil. ^1^H NMR (500 MHz, as a mixture of rotamers) δ 7.72-7.24 (comp, 12 H), 5.61 (br s, 0.5 H), 5.49 (br s, 0.5 H), 5.27-5.07 (comp, 2 H), 3.96-3.81 (m, 1 H), 3.39-3.24 (m, 1 H), 2.61-2.43 (m, 1 H), 2.34-2.20 (m, 1 H), 2.12-2.00 (m, 1 H), 1.94 (d, *J* = 11.0 Hz, 1 H), 1.74-1.59 (m, 1 H). ^13^C NMR (126 MHz, as a mixture of rotamers) δ 155.0, 154.8, 146.5, 144.6, 142.1, 141.9, 139.0, 136.9, 136.8, 129.2 (q, *J*_C−F_ = 31.5 Hz), 128.4, 127.9, 127.8, 127.3, 125.6 (q, *J*_C−F_ = 3.8 Hz), 123.3 (q, *J*_C−F_ = 272.2 Hz), 123.2, 122.7, 122.5, 67.0, 57.6, 57.3, 43.6, 39.5, 38.6, 30.2. HRMS (ESI) *m/z* calcd for C_26_H_22_F_3_NNaO_2_ (M+Na)^+^, 460.1495; found 460.1499.

8*-(4-(Trifluoromethyl)phenyl)-2,3,4,5-tetrahydro-1H-1,5-methanobenzo[c]azepine (****3****).* A mixture of **2** (243 mg, 0.55 mmol), 10% Pd/C (85 mg) and EtOH (4 mL) was stirred under a H_2_ balloon at room temperature for 5 h. The mixture was filtered through a pad of Celite, which was washed with CH_2_Cl_2_ (2 mL), and the combined filtrate and washings were concentrated under reduced pressure. The crude product was purified via flash column chromatography (SiO_2_), eluting with MeOH/NEt_3_/EtOAc (1:1:8), to afford 150 mg (90%) of **3** as a colorless oil. ^1^H NMR (500 MHz) δ 7.76 (s, 1 H), 7.68 (q, *J* = 8.5 Hz, 4 H), 7.56 (dd, *J* = 7.5, 1.5 Hz, 1 H), 7.34 (d, *J* = 8.0 Hz, 1 H), 7.11 (br s, 1 H), 4.67 (d, *J* = 2.5 Hz, 1 H), 3.38 (s, 1 H), 3.07 (dd, *J* = 13.0, 5.5 Hz, 1 H), 2.49 (td, *J* = 12.5, 5.0 Hz, 1 H), 2.42 – 2.33 (comp, 2 H), 2.22 (td, *J* = 12.5, 5.0 Hz, 1 H), 1.67 (d, *J* = 13.5 Hz, 1 H). ^13^C NMR (126 MHz) δ 146.1, 144.2, 139.6, 139.1, 129.3 (q, *J*_C−F_ = 32.5 Hz), 128.6, 127.4, 125.7 (q, *J*_C−F_ = 3.8 Hz), 124.2 (q, *J*_C−F_ = 272.2 Hz), 123.2, 123.1, 58.0, 42.7, 38.8, 38.1, 28.3. HRMS (ESI) *m/z* calcd for C_18_H_17_F_3_N (M+H)^+^, 304.1308; found 304.1309.

*3-((1S,5R)-8-(4-(Trifluoromethyl)phenyl)-1,3,4,5-tetrahydro-2H-1,5-methanobenzo[c]azepin-2-yl)propan-1-ol (FEM-1689).* To a solution of **3** (191.2 mg, 0.63 mmol) in acetonitrile (6 mL) was added K_2_CO_3_ (261.2 mg, 1.89 mmol), followed by 3-bromopropan-1-ol (175.2 mg, 1.26 mmol). The reaction mixture was heated to 60 °C for 21 h. The reaction mixture was cooled to room temperature and filtered, and the filtrate was concentrated under reduced pressure. The residue was suspended in 1 M aqueous HCl (5 mL) and washed with ether (5 mL). The aqueous phase was basified (pH ~ 8) with 2 M aqueous NaOH and extracted with CH_2_Cl_2_ (3 × 20 mL). The extracts were combined, dried (Na_2_SO_4_), filtered, and concentrated reduced pressure. The residue was purified by column chromatography (SiO_2_) using CH_2_Cl_2_:MeOH (30:1 to 15:1) as eluant to yield 115.8 mg (51%) FEM-1689 as a colorless oil. LCMS, retention time 5.31 min, 97% pure.^1^H NMR (500 MHz) δ 7.69 (s, 4 H), 7.49 (dd, *J* = 7.6, 1.7 Hz, 1 H), 7.39 (s, 1 H), 7.31 (d, *J* = 7.6 Hz, 1 H), 4.16 (d, *J* = 4.6 Hz, 1 H), 3.93 – 3.79 (comp, 2 H), 3.21 (br. s, 1 H), 2.84 (dd, *J* = 11.7, 5.7 Hz, 1 H), 2.77 (ddd, *J* = 12.0, 7.5, 3.8 Hz, 1 H), 2.39 (ddd, *J* = 12.2, 7.8, 3.8 Hz, 1 H), 2.32 – 2.26 (m, 1 H), 2.07 – 1.98 (m, 1 H), 1.97 (d, *J* = 11.0 Hz, 1 H), 1.86 – 1.76 (m, 1 H), 1.75 – 1.66 (m, 1 H), 1.61 – 1.55 (m, 1 H), 1.50 (td, *J* = 11.9, 4.8 Hz, 1 H). ^13^C NMR (126 MHz) δ 146.9, 145.1, 139.5, 138.4, 129.30 (q, *J* _C-F_ = 32.4 Hz), 127.5, 127.4, 125.8 (q, *J* _C-F_ = 3.8 Hz), 124.6 (q, *J* _C-F_ = 272.4 Hz), 123.2, 123.1, 64.9, 63.6, 56.7, 47.1, 44.7, 39.7, 30.1, 27.2. HRMS (FIA) *m/z* calcd for C_21_H_23_F_3_NO (M+H)^+^ 362.1726; found 362.1728.

*Glide Docking Calculations.* Molecular docking into σ_2_R/TMEM97 was performed using the Glide module with standard precision (SP) in Maestro of the Schrodinger software suite (release 2023-1) (9). We have shown that the 1*S,*5*R*-enantiomers of several piperazine-substituted norbenzomorphans have higher affinities for σ_2_R/TMEM97 than their 1*R,*5*S*-enantiomers (10), and the known structures of σ_2_R/TMEM97 complexed with amine ligands uniformly show the amino groups are protonated (4). Accordingly, LigPrep was used to generate starting ligand structures of the 1*S,*5*R*-enantiomers of SAS-0132 and FEM-1689, and the structures in which the more basic amino group of the ligand is protonated were used. The published structure (2.71 Å; pdb accession 7M94) of bovine σ_2_R/TMEM97 complexed with roluperidone has four protomers in the asymmetric unit (4), but Chain C was used for docking studies. All lipids, ions, and waters were removed prior to grid preparation, leaving only the protein and ligand. Hydrogen atoms were added, and the protein was further refined by assigning H-bonds and minimizing energy for the OPLS4 force field. After protein preparation, Asp29 and Glu73 were deprotonated, whereas Tyr150 was protonated. The grid used for docking was centered on the location of the co-crystallized ligand roluperidone, and was 20 Å in the x, y, and z dimensions. During docking computations, a constraint requiring an interaction between a protonated amino group on the ligand and Asp29 was applied because such H-bonds and salt-bridges are conserved in known structures of complexes of σ_2_R/TMEM97 with basic amines. Poses were ranked by docking score.

**Other Supp Materials:**

**Dose Response Curves for Receptor Binding for FEM-1689 (PDSP #52656)**

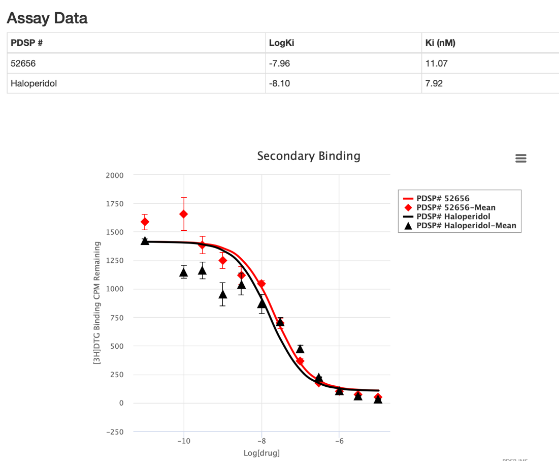

TMEM97

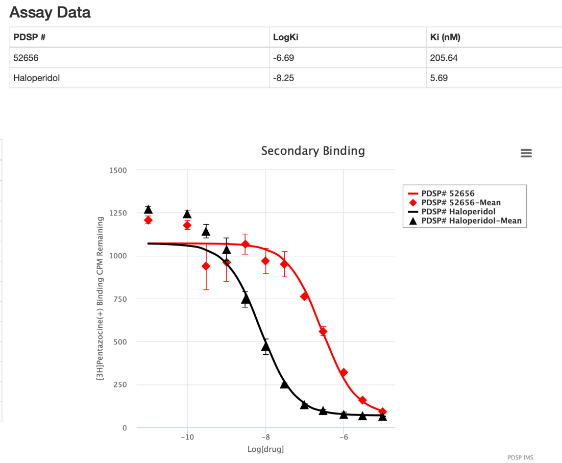

Sigma 1 Receptor

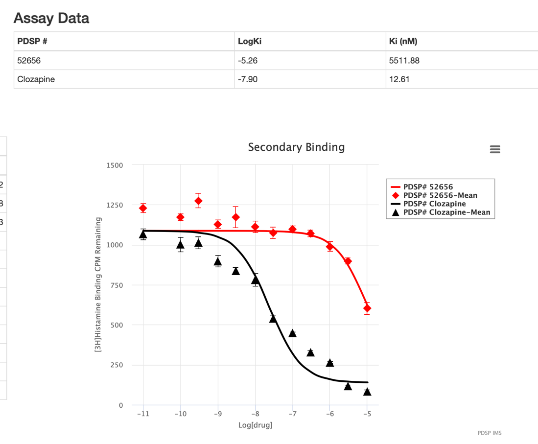

H4

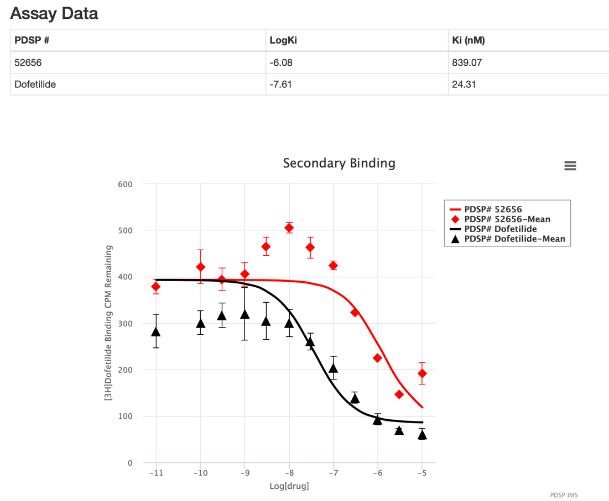

HERG

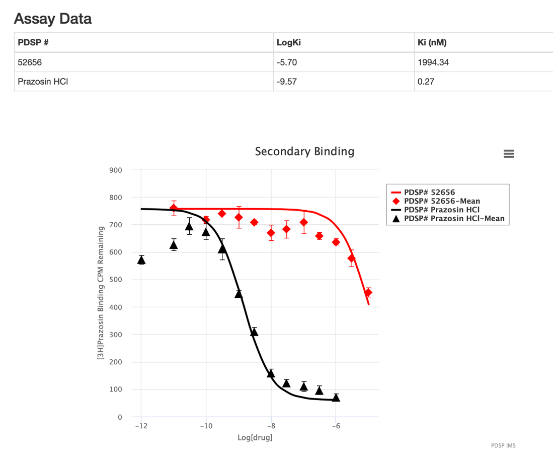

Alpha1A

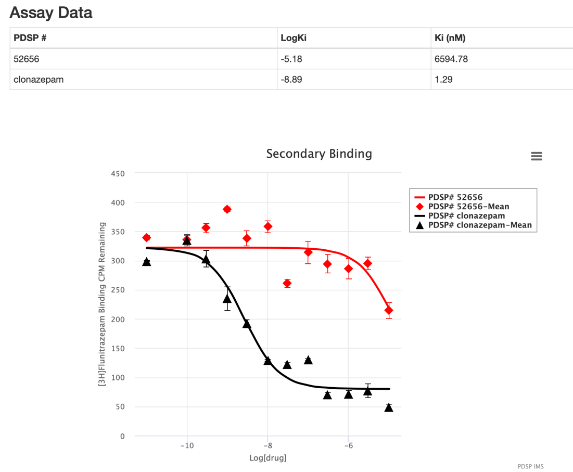

BZP Rat Brain

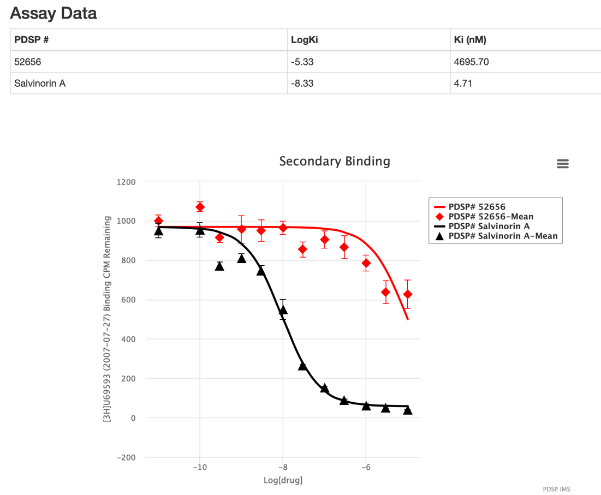

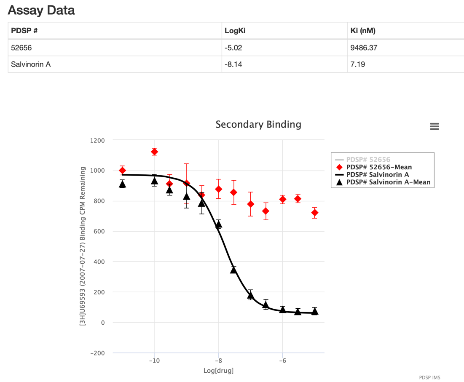

KOR

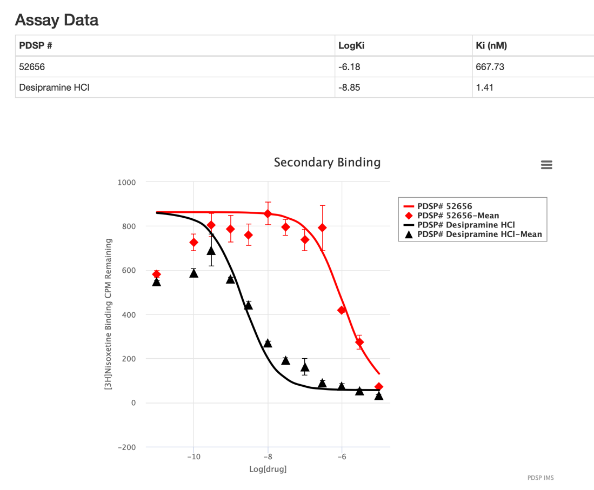

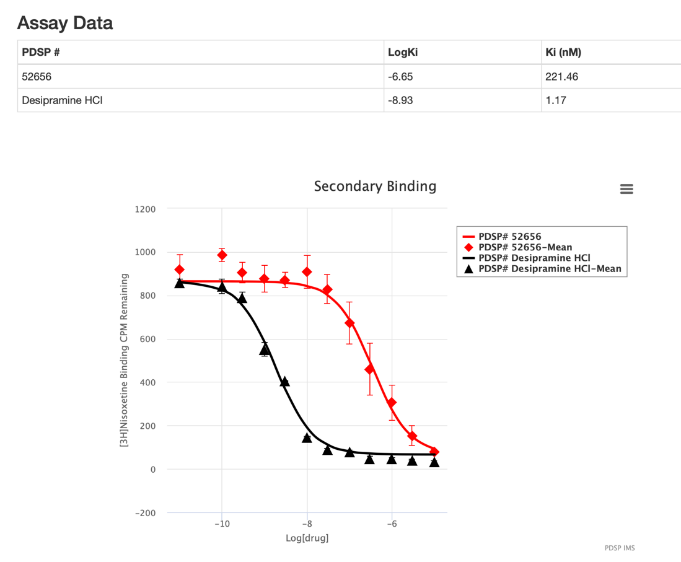

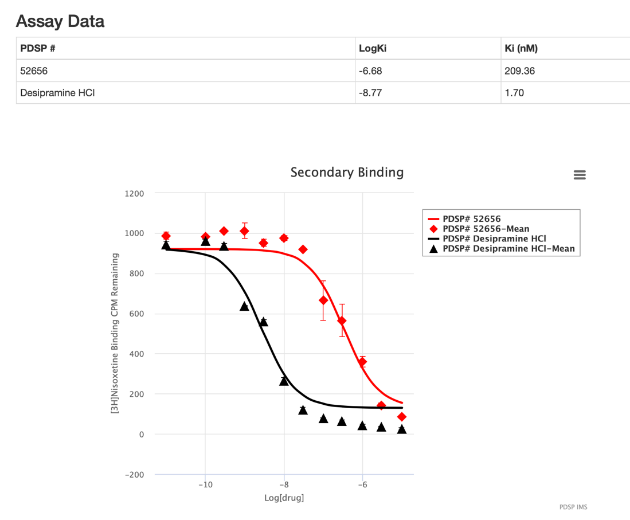

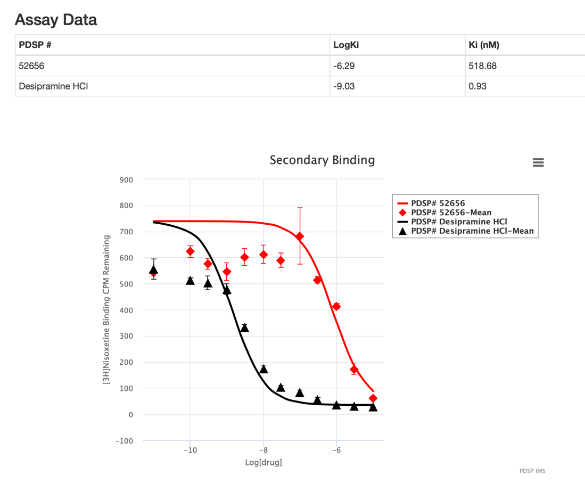

NET

**NMR Data for FEM-1689**

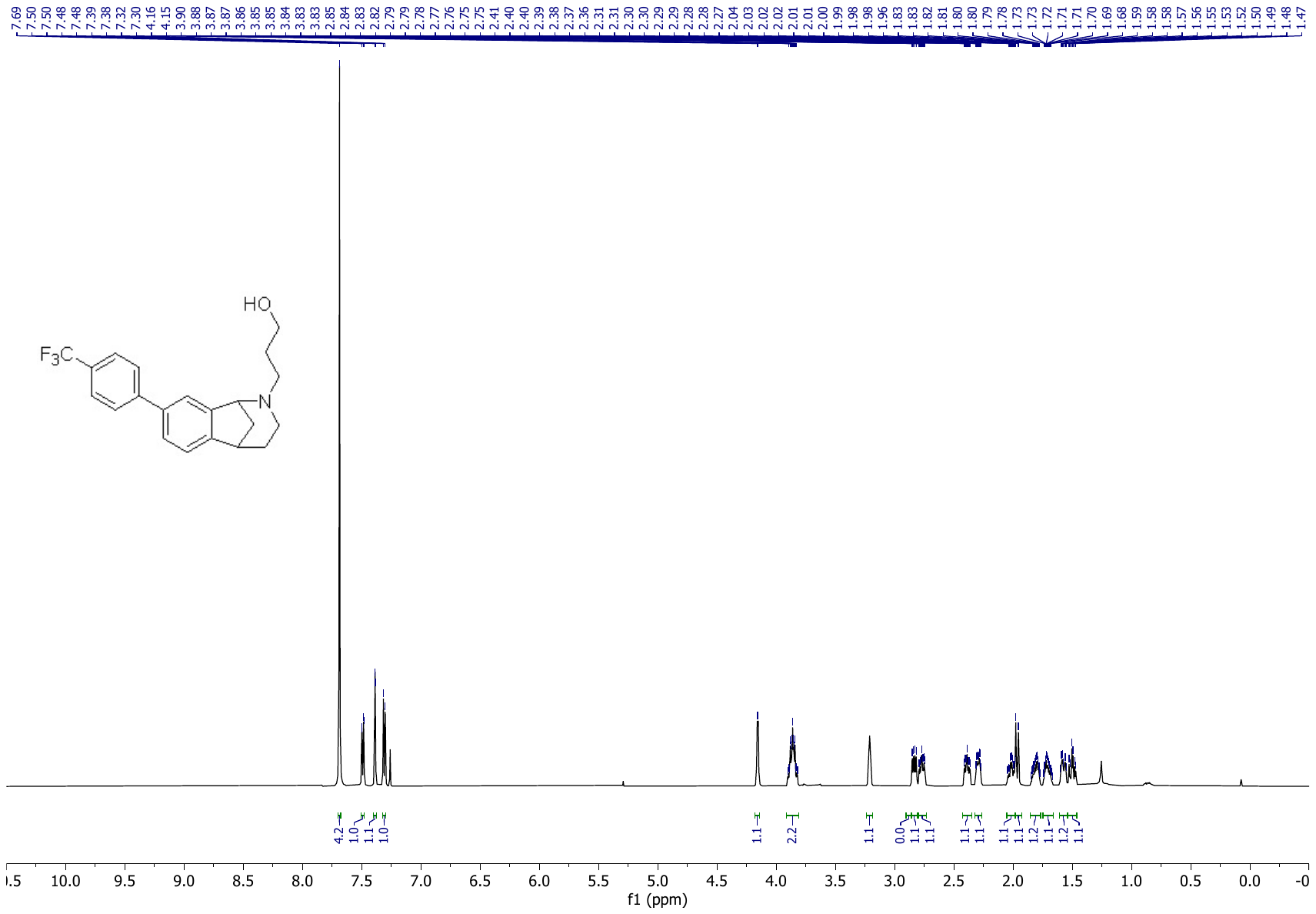

^1^H NMR spectrum in CDCl_3_

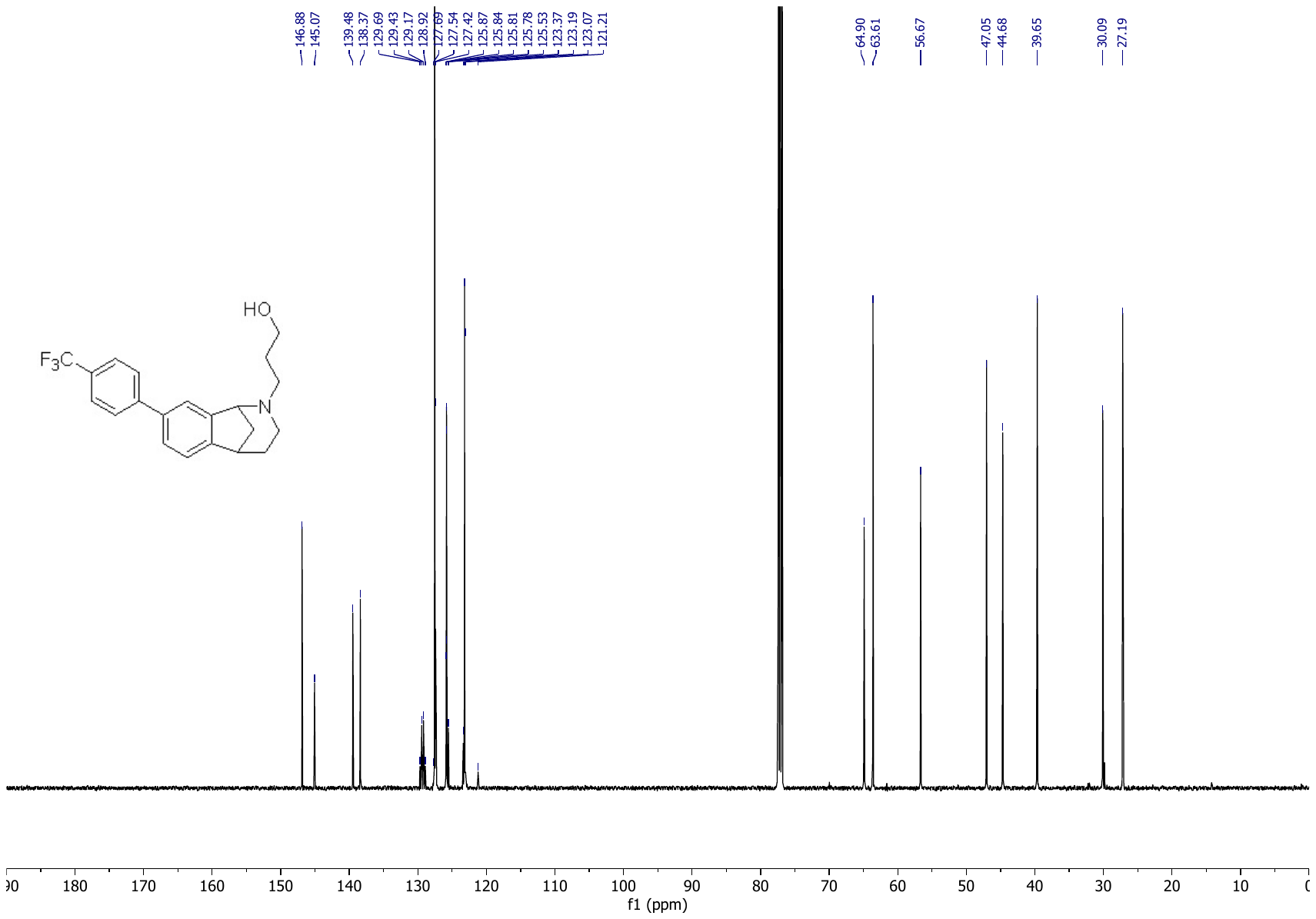

^13^C NMR spectrum in CDCl_3_

**LCMS data for FEM-1689**

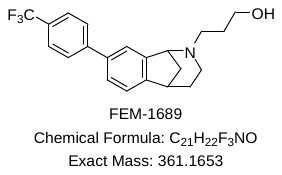

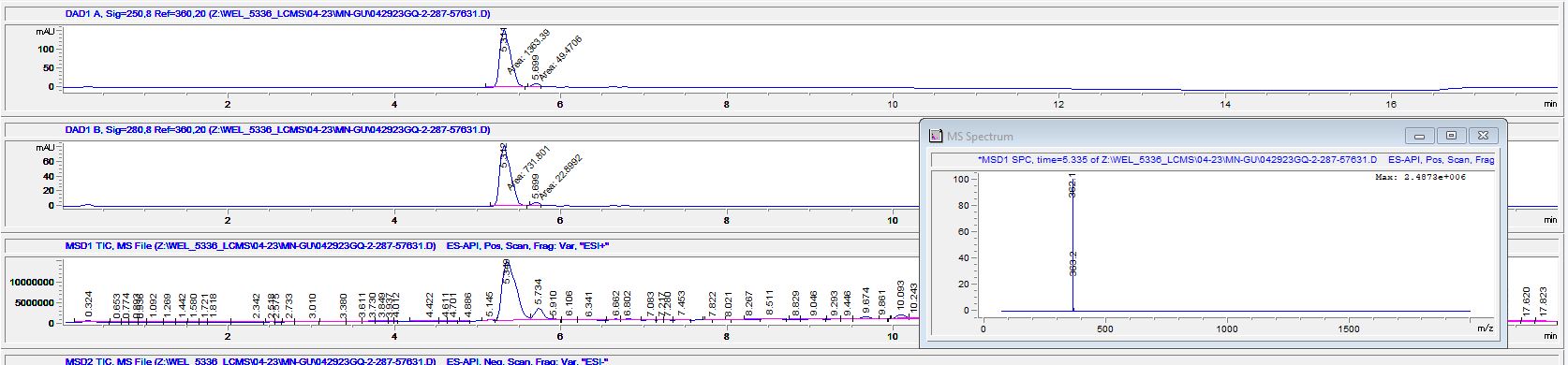

Retention Time, 5.31 min, purity 96.5
